# Photoinduced Isomerization Sampling of Retinal in Bacteriorhodopsin

**DOI:** 10.1101/2021.09.16.460656

**Authors:** Zhong Ren

## Abstract

Photoisomerization of retinoids inside a confined protein pocket represents a critical chemical event in many important biological processes from animal vision, non-visual light effects, to bacterial light sensing and harvesting. Light driven proton pumping in bacteriorhodopsin entails exquisite electronic and conformational reconfigurations during its photocycle. However, it has been a major challenge to delineate transient molecular events preceding and following the photoisomerization of the retinal from noisy electron density maps when varying populations of intermediates coexist and evolve as a function of time. Here I report several distinct early photoproducts deconvoluted from the recently observed mixtures in time-resolved serial crystallography. This deconvolution substantially improves the quality of the electron density maps hence demonstrates that the all-*trans* retinal undergoes extensive isomerization sampling before it proceeds to the productive 13-*cis* configuration. Upon light absorption, the chromophore attempts to perform *trans*-to-*cis* isomerization at every double bond together with the stalled *anti*-to-*syn* rotations at multiple single bonds along its polyene chain. Such isomerization sampling pushes all seven transmembrane helices to bend outward, resulting in a transient expansion of the retinal binding pocket, and later, a contraction due to recoiling. These ultrafast responses observed at the atomic resolution support that the productive photoreaction in bacteriorhodopsin is initiated by light-induced charge separation in the prosthetic chromophore yet governed by stereoselectivity of its protein pocket. The method of a numerical resolution of concurrent events from mixed observations is also generally applicable.

**Significance Statement:** Photoisomerization of retinal is a critical rearrangement reaction in many important biological processes from animal vision, non-visual light effects, to bacterial light sensing and harvesting. It has been a major challenge to visualize rapid molecular events preceding and following photoisomerization so that many protein functions depending on such reaction remain vaguely understood. Here I report a direct observation of the stereoselectivity of bacteriorhodopsin hence delineate the structural mechanism of isomerization. Upon a light-induced charge separation, the retinal in a flat conformation attempts to perform double bond isomerization and single bond rotation everywhere along its polyene chain before it proceeds to the specific, productive configuration. This observation improves our understanding on how a non-specific attraction force could drive a specific isomerization.

## Introduction

Bacteriorhodopsin (bR) pumps protons outward from the cytoplasm against the concentration gradient via photoisomerization of its retinal chromophore. The trimeric bR on the native purple membrane shares the seven transmembrane helical fold and the same prosthetic group (Fig. S1) with large families of microbial and animal rhodopsins (Ernst et al., 2014; Kandori, 2015). An all-*trans* retinal in the resting state is covalently linked to Lys216 of helix G through a protonated Schiff base (SB; Fig. S1e), of which the double bond C_15_=N_ζ_ is also in *trans* (traditionally also noted as *anti* (McCarty, 1970)). Upon absorption of a visible photon, the all-*trans* retinal in bR isomerizes efficiently and selectively to adopt the 13-*cis* configuration (Govindjee et al., 1990; Tittor and Oesterhelt, 1990). In contrast, an all-*trans* free retinal in organic solvents could isomerize about various double bonds with a preference to form 11-*cis*, but much slower (Hamm et al., 1996; Logunov et al., 1996) and with poor quantum yields (Freedman and Becker, 1986; Koyama et al., 1991).

A broad consensus is that the isomerization event takes place around 450-500 fs during the transition from a blue-shifted species I to form a red-shifted intermediate J (Herbst, 2002; Mathies et al., 1988). This ultrafast event is explained in excited state dynamics as a barrierless reaction, or at least with a very small activation barrier, from the Frank-Condon point to the conical intersection between the potential energy surfaces of the ground and excited states (Birge, 1990; Gozem et al., 2017). Various molecular events prior to the isomerization have also been detected. Vibrational spectroscopy showed a variety of possible motions, such as torsions about C_13_=C_14_ and C_15_=N_ζ_, H-out-of-plane wagging at C_14_, and even protein responses (Diller et al., 1995; Kobayashi et al., 2001). Nevertheless, the species I or a collection of species detected before 30 fs remain in a good *trans* configuration about C_13_=C_14_ instead of a near 90° configuration (Zhong et al., 1996). Recently, deep-UV stimulated Raman spectroscopy revealed strong signals of Trp and Tyr motions in the protein throughout the I and J intermediates (Tahara et al., 2019). Despite extensive studies, fundamental questions on the photoisomerization of retinal remain unanswered at the atomic resolution. It is assumed that the specific isomerization at C_13_=C_14_ in bR occurs faster with a high quantum yield compared to the slower, nonspecific isomerization at multiple sites for free retinal in solution and gas phase (Freedman and Becker, 1986; Hamm et al., 1996; Kiefer et al., 2019; Koyama et al., 1991; Logunov et al., 1996) because the interactions between the retinal chromophore and its protein environment somehow catalyze the photoreaction by tuning the potential energy surfaces of the ground and/or excited states. What is the structural basis of such tuning or catalysis? What is the quantum mechanical force that causes the all-*trans* retinal to isomerize specifically to 13-*cis* in bR? Why not isomerize elsewhere in bR? How is the quantum yield of this specific isomerization enhanced by the protein compared to those of free retinal in solution? Does any isomerization sampling occur, that is, is isomerization even attempted at other double bonds? This work addresses these questions by solving a series of structures of the early intermediates based on the electron density maps unscrambled from the published serial crystallography datasets using singular value decomposition (SVD). These structures of “pure” photoproducts at atomic resolution reveal widespread conformational changes in all seven helices prior to the all-*trans* to 13-*cis* isomerization and after its completion, suggesting that isomerization sampling takes place in bR, where rapid photoisomerizations and single bond rotations are attempted everywhere along the polyene chain of the retinal before the only successful one flips the SB at ~500 fs. The implication of these findings to the proton pumping and directional conductance is presented in a companion paper (Ren, 2021).

Several international consortia carried out large operations of serial crystallography at X-ray free electron lasers (XFELs). It is now possible to capture transient structural species at room temperature in the bR photocycle as short-lived as fs (Brändén and Neutze, 2021). Compared to cryo-trapping, authentic structural signals from these XFEL data are expected to be greater in both amplitude and scope. However, the signals reported so far do not appear to surpass those obtained by cryo-trapping methods, suggesting much needed improvements in experimental protocols and data analysis methods. Two major sources of data are used in this study (Table S1). Nogly et al. captured retinal isomerization to 13-*cis* by the time of 10 ps and attributed its specificity to the H-bond breaking between the SB and a water (Nogly et al., 2018). Kovacs et al. deposited datasets at many short time delays (Kovacs et al., 2019). Those sub-ps datasets demonstrate oscillatory signals at frequencies around 100 cm^-1^ (see below). But no signal of conformational change, other than the extensive oscillations, is found from these data in the transmembrane helices or the chromophore depicting the retinal isomerization despite the original report. The essence of this work is a numerical resolution of structural heterogeneity, a common difficulty often encountered in cryo trapping and time-resolved crystallography. To what extend a specific structural species can be enriched in crystals depends on the reaction kinetics governed by many experimental parameters including but not limited to the fluence, wavelength, and temperature of the light illumination. While it is possible to reach higher fractional concentrations at specific time points for more stable species such as K and M states of bR due to the ratio between the rates going into and exiting from that species, transient species such as I and J are often poorly populated. If such structural heterogeneity is not resolved, it is very difficult, if not impossible, to interpret the electron density maps and to refine the intermediate structures (Ren et al., 2013). An assumption in nearly all previous studies has been that each dataset, at a cryo temperature or at a time delay, is derived from a mixture of a single conformer of a photoinduced species and the ground state. Therefore, the difference map reveals a pure intermediate structure. This assumption is far from the reality thus often leads to misinterpretation of the observed electron density maps. This work is yet another case study to demonstrate the application of our analytical protocol based on SVD (Methods) that makes no assumption on how many excited intermediates that contribute to the captured signals at each time point (Ren, 2019; Ren et al., 2013; Yang et al., 2011). More importantly, this work showcases that our resolution of structural heterogeneity enables new mechanistic insights into the highly dynamic chemical or biochemical processes.

## Results and Discussion

A total of 22 datasets and 18 time points up to 10 ps are analyzed in this study (Table S1). Difference Fourier maps at different time points and with respect to their corresponding dark datasets are calculated according to the protocols previously described (Methods). A collection of 126 difference maps at short delays ≤ 10 ps are subjected to singular value decomposition (SVD; Methods) followed by a numerical deconvolution using the previously established Ren rotation in a multi-dimensional Euclidean space (Ren, 2016, 2019). Such resolution of electron density changes from mixed photoexcited species in the time-resolved datasets results in four distinct intermediate structures in the early photocycle (Fig. 1), which are then refined against the reconstituted structure factor amplitudes (Table S2; Methods).

**Figure 1.**
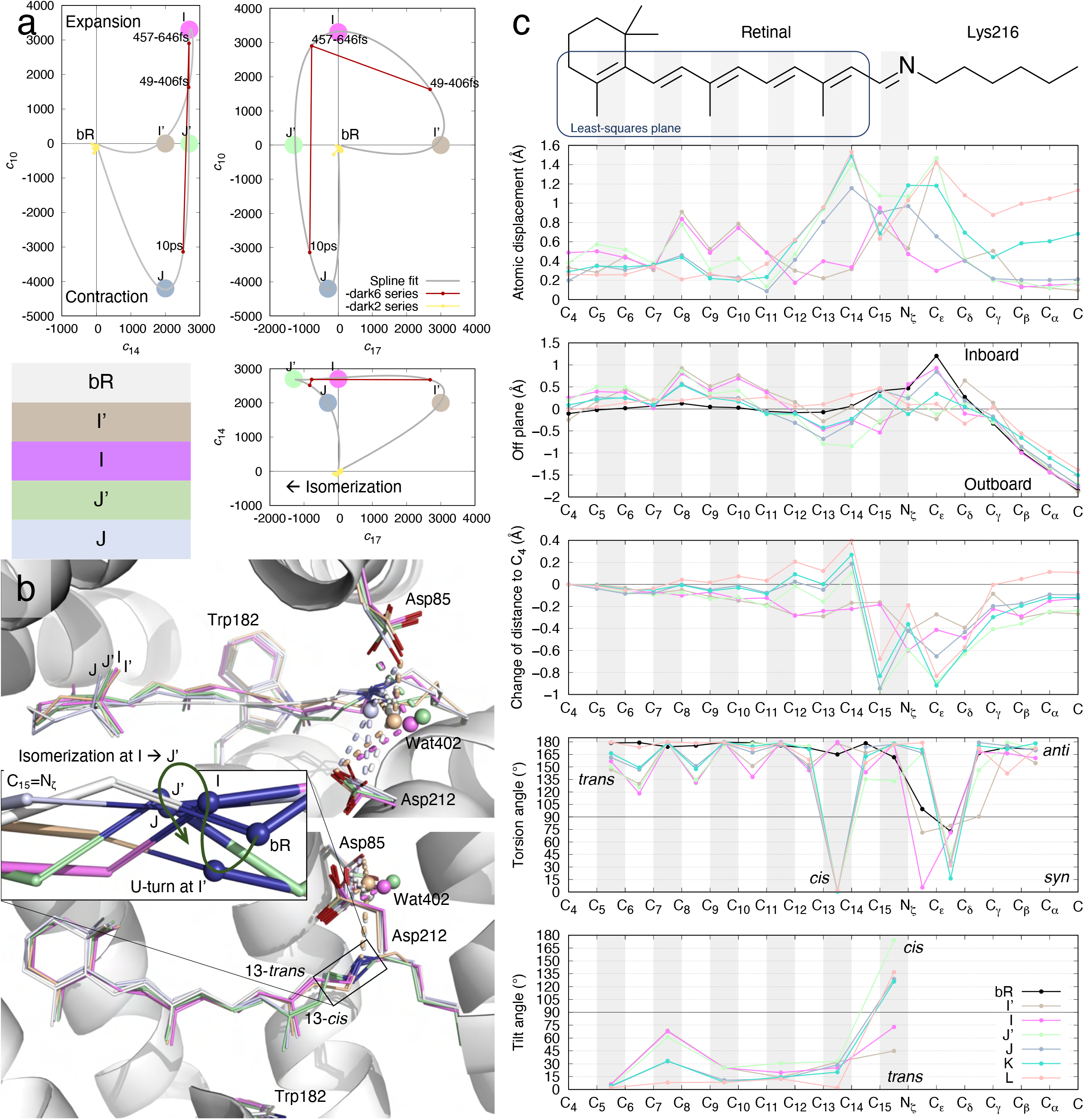
Intermediate species in early photocycle identified in SVD space. (a) Multi-dimensional spaces of SVD. The SVD analysis of difference Fourier maps at short delays ≤ 10 ps results in time-dependent coefficients *c_k_*(*t*), where *k* = 1, 2, …, each corresponding to a time-independent components ***U***_*k*_. Each raw difference map at a time delay *t* can be closely represented by a linear combination of these components, *c*_1_(*t*)***U***_1_ + *c*_2_(*t*)***U***_2_ + …, that is called a reconstituted difference map. Each of these components **U**_*k*_ and the reconstituted difference maps can be rendered in the same way as an observed difference map. The coefficient set *c*_*k*_(*t*) is therefore a trace of the photocycle trajectory, when these time-dependent functions are plotted in a multi-dimensional space or plotted together against the common variable of time *t*. Coefficients corresponding to components ***U***_10_, ***U***_14_, and ***U***_17_ are plotted in three orthographical views. Three time points from Nogly et al. in small red dots contain ***U***_14_ equally. These time points vary in ***U***_10_ and ***U***_17_. Datasets from Kovacs et al. in small yellow dots do not carry any of these signals, therefore cluster near the origin. The component map of ***U***_10_ is displayed in Fig. 3c. ***U***_14_ is displayed in Figs. 2a and S5. ***U***_17_ is displayed in Fig. 2b. Several apices of the spline fitting are chosen as the potential pure states of I’, I, J’, and J marked by large dots. This choice is only an approximate due to the insufficient number of time points observed. (b) The refined retinal conformations compared with the resting state in white. I’, I, J’, and J are in beige, purple, green, and bluish gray, respectively. The creased S-shape of the retinal is established in I’ (Fig. 2) and easing gradually (Fig. 1bc). 13-*trans* in I’ and I, 13-*cis* in J’ and J are two discrete conformations without any other conformations in the middle (c 4^th^ panel). The inset is a zoom-in view of the SB double bond C_15_=N_ζ_. The SB N_ζ_ atom performs two rapid U- turns at ~30 fs and ~500 fs. (c) Conformational parameters calculated from the refined chromophore. The chemical structure of the chromophore on top is aligned to the horizontal axis. Double bonds are shaded in gray. Intermediates K and L are taken from the companion paper (Ren, 2021). 1) Atomic displacements of each intermediate from the resting state show greater changes in the proximal segment around the SB (top panel). See Fig. S1 for definitions of proximal, distal, inboard, outboard, and other orientations in bR molecule. 2) A plane is least-squares fitted to C_4_ through C_14_ of the resting state as boxed on the chemical structure. The distances of all atoms to this plane in the inboard and outboard directions show the curvature of the chromophore. The creased retinal in early intermediates and the inboard protruding corner at C_ε_ in the resting state are clearly shown (2^nd^ panel). 3) Distances to atom C_4_ are calculated for all refined chromophores. Changes in these distances with respect to the resting state show the shortened chromophore in I’ and I. Once isomerization to 13-*cis* occurs, the segment from C_15_ to C_δ_ around the SB becomes significantly closer to the β-ionone ring due to the Coulombic attraction force, while the distal segment of the retinal from C_14_ and beyond stretches (3^rd^ panel). 4) The torsion angles of single and double bonds quantify *anti/syn* or *trans/cis* for the ground state and all intermediates (4^th^ panel). 5) Only a single bond can be twisted with its torsion angle near 90°. A twisted double bond would be energetically costly. Each double bond is least-squares fitted with a plane. The interplanar angle between a double bond and the corresponding one in the ground state measures the local tilting of the retinal (bottom panel).

### Low frequency oscillations upon photoexcitation

Ten out of 17 major components derived from the sub-ps delays of Kovacs et al. (Fig. S2c) describe five two-dimensional oscillatory behaviors at frequencies ranging from 60 to 400 cm^-1^ (Fig. S3). Compared to a bond stretching frequency commonly observed in vibrational spectroscopy, these oscillations are at much lower frequencies. The lowest frequency is 61±2 cm^-1^, that is, a period of 550±20 fs (Fig. S3a), observed for more than an entire period, which matches exactly the oscillation detected in transient absorption changes in visual rhodopsin (Wang et al., 1994). The second lowest frequency at 150±3 cm^-1^ is observed for more than one and a half periods, and the highest frequency oscillation is observed for nearly six periods at 396±3 cm^-1^ (Fig. S3). Although these ten components follow the oscillatory time dependencies nicely, they do not show any association with the chromophore or any secondary structure of the protein (Fig. S4). Similar oscillatory components were also extracted from the time-resolved XFEL datasets of carbonmonoxy myoglobin (Ren, 2019). Therefore, the same conclusion stands, that is, these low frequency vibrations induced by short laser pulses often detected by ultrafast spectroscopy are the intrinsic property of a solvated protein molecule (Johnson et al., 2014; Liebel et al., 2014). However, the functional relevance of these oscillations is unclear. Interestingly, the isomerization sampling and productive photoisomerization observed in this study occur within the first oscillatory period at the lowest frequency. While such coincidence begs the question whether the protein oscillation is required for isomerization (see below), direct evidence is lacking in these XFEL data to support any functional relevance of these oscillatory signals.

These oscillations originate solely from the sub-ps datasets (Fig. S2ab yellow, green, blue, and purple traces) of Kovacs et al. derived from excessive peak power density of their pump laser (Kovacs et al., 2019; Miller et al., 2020). The datasets of Nogly et al. in the same delay range carry coefficients ≈ 0 corresponding to these oscillatory components, that is, free of oscillation (Fig. S2ab red and brown traces near the origin). On the other hand, no dataset of Kovacs et al. features significantly nonzero coefficient other than those corresponding the oscillatory components (Fig. 1a yellow trace near the origin). Therefore, these datasets containing oscillations do not carry any interpretable signal in the protein or on the chromophore, at least not in the outstanding components identified by SVD (Fig. S2c). As Miller et al. pointed out, it is entirely possible that the specimens were pumped onto higher potential energy surfaces due to multiphoton absorption and the subsequent decays follow some biologically irrelevant reaction pathways (Miller et al., 2020).

### Intermediates I’, I, and expansion of retinal binding pocket

In contrast to the oscillating signals, three components ***U***_10_, ***U***_14_, and ***U***_17_ reveal strong light-induced structural signals in terms of both extensiveness and quality (Figs. 2ab and S5). These signals originate exclusively from a few time points of Nogly et al., too few to fit the time dependency with exponentials. Instead, a spline fitting through these time points gives rise to the estimated coefficients *c*_10_, *c*_14_, and *c*_17_ in the linear combination of *c*_10_***U***_10_ + *c*_14_***U***_14_ + *c*_17_***U***_17_ for reconstructing the electron density maps of the states I, J, and their respective precursors I’, J’ (Fig. 1a). A reconstituted difference map of I’ – bR = 2000***U***_1_4 + 3000***U***_17_ (Fig. 2c) is located on the spline trajectory from the origin, that is, bR at the time point of 0-, to the first time point of 49-406 fs (PDB entry 6g7i). This state is denoted I’ as a precursor leading to the I state judged by the time point at ~30 fs. However, this is not to say that a single species I’ exists around 30 fs. Quite the opposite, the population of the time-independent conformational species I’ rises and falls and peaks around 30 fs, while many other species during isomerization sampling coexist with I’ at the same time (see below). The reconstituted difference map is used to calculate a set of structure factor amplitudes that would produce this difference map of I’ – bR (Methods). And the structure of I’ is refined against this reconstituted dataset (beige; Figs. 2cd and S6). The same protocol is used to refine the structure of I state (purple; Fig. S7) with a reconstituted difference map I – bR = 3300***U***_10_ + 2700***U***_14_ (Fig. 3d). This SVD-dependent refinement strategy extends the commonly used method based on an extrapolated map to another level. This newly developed method is able to refine a structure against any linear combination of signal components while eliminating noise and systematic error components, and components identified as other intermediate species mixed in the data. Therefore, this method enables the refinement of an unscrambled, hence pure, structural species (Methods).

**Figure 2.**
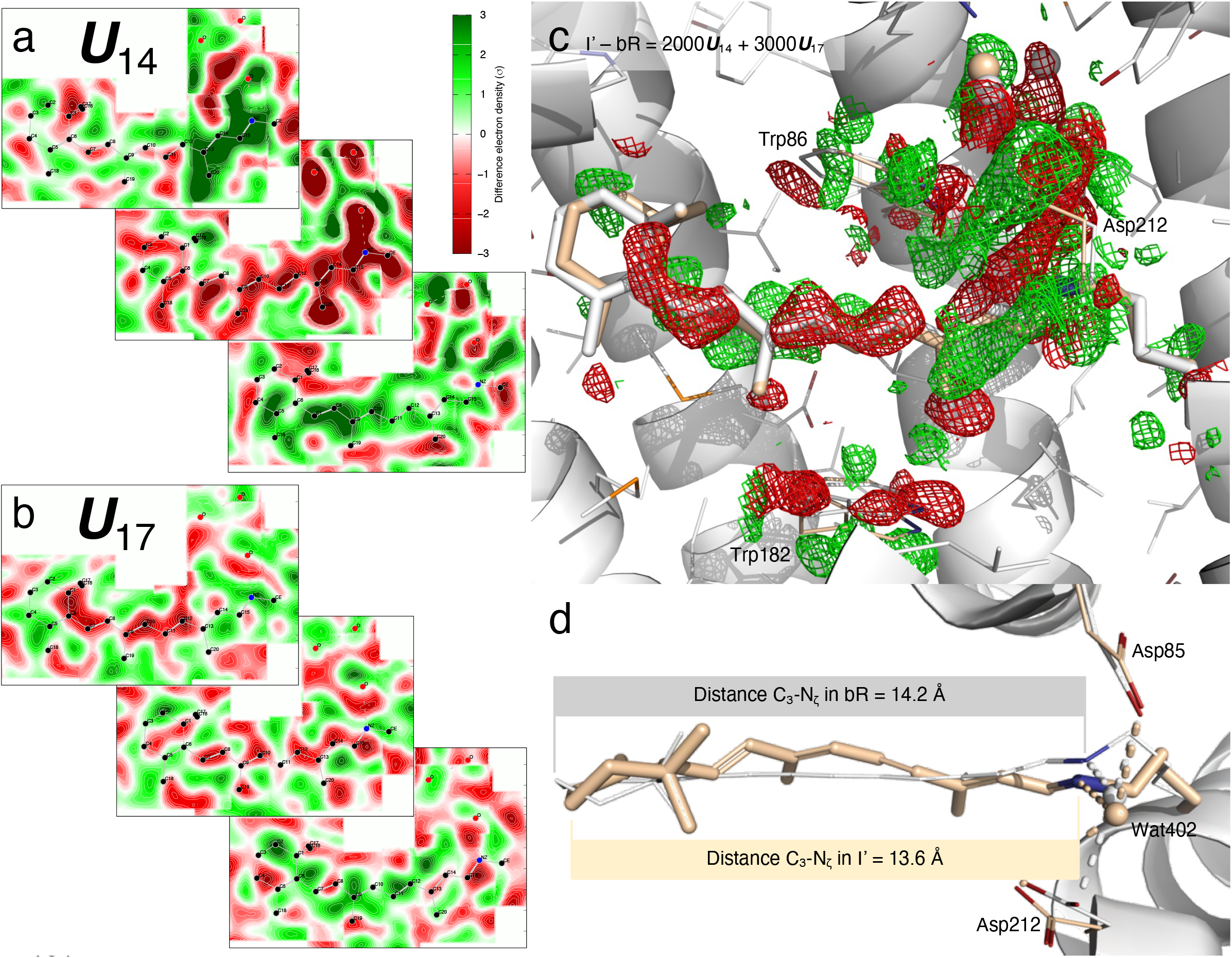
Shortened retinal in S-shape since earliest intermediate I’. (a) Cross sections of component map ***U***_14_. The middle cross section is an integration ±0.2 Å around the curved surface through the retinal. The top cross section is an integration 1.2-1.8 Å outboard from the retinal surface and the bottom one is an integration 0.8-1.2 Å inboard. See Fig. S1 for definitions of inboard, outboard, proximal, distal, and other orientations in bR molecule. Green and red indicate electron density gain and loss, respectively. Nearly the entire retinal is in negative densities. The proximal segment and three waters are in intense negative densities. On the other hand, strong positive densities flank the proximal and distal segments from the outboard and inboard, respectively. Such signal distribution results in the S-shaped retinal by the refinement shown in (d). (b) Cross sections of component map ***U***_17_. The middle cross section is an integration ±0.2 Å around the surface through the retinal. The top panel is an integration 0.5-0.9 Å outboard and the bottom is an integration 0.8-1.2 Å inboard. Negative and positive densities flank the retinal from the outboard and inboard, respectively. (c) Difference map of I’ – bR reconstituted from ***U***_14_ and ***U***_17_ (a and b). The map is contoured at ±3*σ* in green and red mesh, respectively. The opposite displacements of the distal and proximal segments of the retinal are obvious. Extensive signals indicate changes in the water network and Asp85 and 212. (d) Refined retinal conformation in beige overlaid on the resting state in white. This view is orthographical to (c). The marked distances from C_3_ to N_ζ_ show a shortened retinal creased into an S- shape. C_20_ methyl group is tilted 33° toward outboard from its resting state bR (Fig. 1c bottom panel). Wat402 remains in H-bonds with both Asp85 and 212.

**Figure 3.**
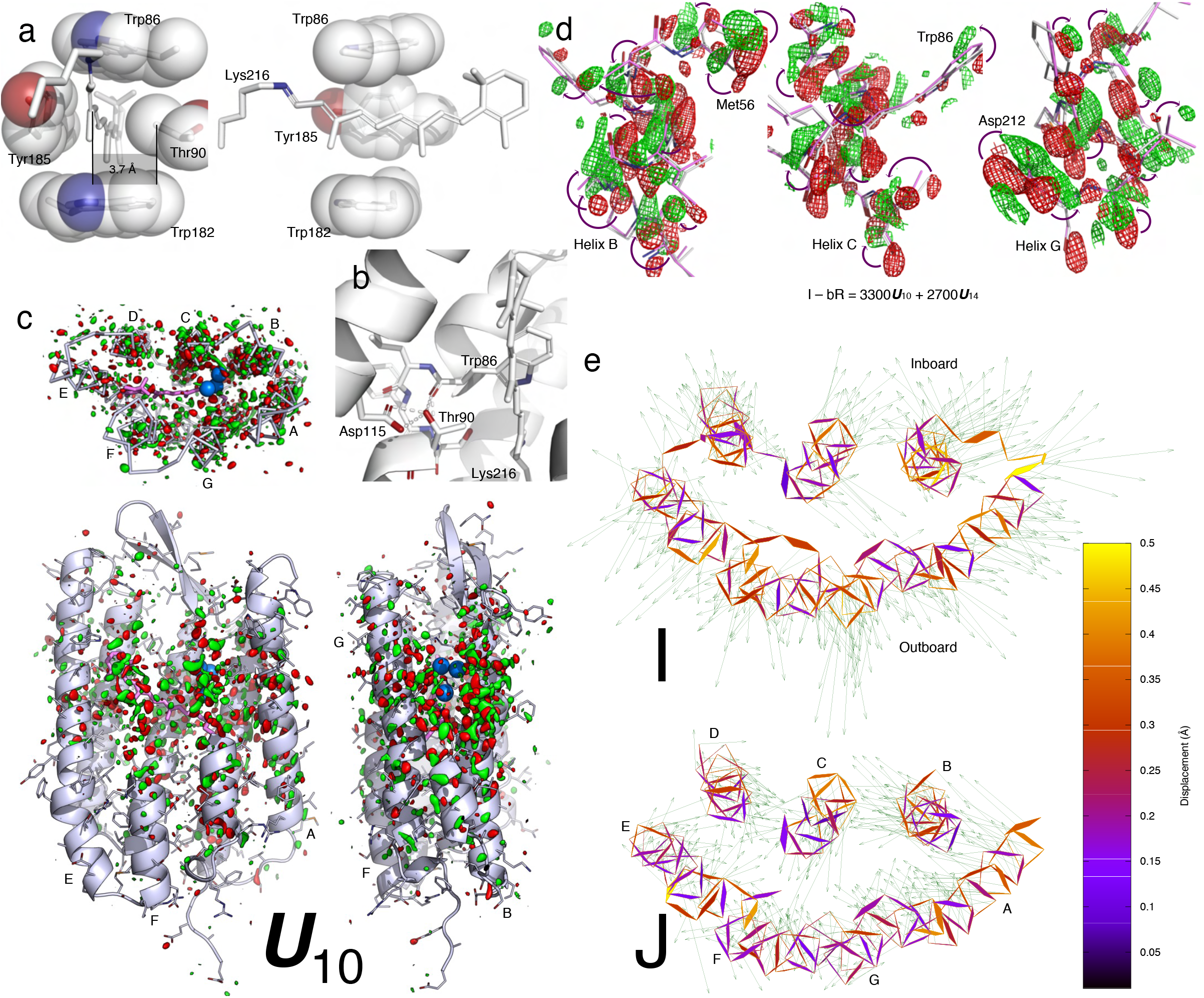
Retinal binding pocket expansion and contraction. (a) Two orthographical views of the retinal tightly boxed at its middle segment. The closest contact is Thr90 and Tyr185 on the inboard and outboard sides of the retinal plane, respectively. The minimum distance between them is 7.0 Å = 4*rc* + 0.2 Å, where *rc* = 1.7 Å is the van der Waals radius of C atom. The C_γ_ methyl group is 3.7 Å = 2*rc* + 0.3 Å away from the retinal plane. See also (Kandori, 2015). (b) H-bond network involving Thr90. The O_γ_ hydroxyl group of Thr90 is strongly H-bonded with the main chain carbonyl of Trp86 in helix C and the carboxyl group of Asp115, simultaneously. These H-bonds fixate the side chain of Thr90 so that the C_γ_ methyl group is pointing to the retinal plane directly. (c) Component map **U**_10_. The main chain and side chains of the protein are rendered with ribbons and/or sticks. The retinal and Lys216 are in purple sticks. Several key waters are in blue spheres. Parts of the structure are omitted to reveal more of the interior. The map is contoured at ±3*σ* in green and red, respectively. Three orthographical views of **U**_10_ clearly show that the signals are distributed around the middle segment of the molecule and taper off to both cytoplasmic and extracellular surfaces. The signals also concentrate on all seven helices. (d) Reconstituted difference map I – bR from **U**_10_ (c) and **U**_14_ (Figs. 2a and S5). The map is contoured at ±2.5*σ* in green and red mesh, respectively. The difference map at three middle segments of helices B, C, and G show main chain displacements toward inboard or outboard as indicated by the arrows marking the negative and positive pairs of densities. These difference densities are the direct evidence of the expansion of the retinal binding pocket. The refined structure of I is in purple, and the resting state is in white. (e) The refined structures of I and J compared with the resting state viewed along the trimer three-fold axis from the extracellular side. Atomic displacements in the main chain from bR to I and J are color coded and marked by arrows with lengths 20× of the actual displacements. All seven helices in I move away from the center except a small segment in helix C showing an expansion of the retinal binding pocket (top panel). However, all seven helices in J move closer to one another showing a contraction with respect to the resting state bR (bottom panel). This contraction is much more significant if compared directly with the expanded I state.

Five geometric parameters are calculated along the long-chain chromophore in the refined intermediate species to depict the transient conformation (Fig. 1c): 1) Atomic displacements along the polyene chain and the lysyl side chain of Lys216 with respect to the ground state structure demonstrate the overall amplitude of conformational change of the chromophore (Fig. 1c top panel). 2) The retinal in the ground state is largely flat except C_15_ and the methyl groups of C_16_ and C_17_ (Fig. S1e). Five consecutive double bonds from C_5_ to C_14_ of the retinal are largely coplanar. Atoms along the polyene chain and the lysyl side chain of Lys216 are located on both sides of this retinal plane. The side toward the three-fold axis of the bR trimer is called inboard; the opposite side is outboard (see Fig. S1 for orientations in bR molecule). Distances from the atoms along the long-chain chromophore to the retinal plane in the ground state are calculated for all intermediates (Fig. 1c 2^nd^ panel). This parameter shows the curvature of the refined conformation. 3) Changes in the distances from the atoms along the chromophore to C_4_ show extension or contraction of the chromophore in various intermediates (Fig. 1c 3^rd^ panel). 4) Torsion angles of the single and double bonds along the chromophore clearly demonstrate *syn*/*anti* and *cis*/*trans* conformation as well as possible deviation from the ideal bond geometry in the refined intermediates (Fig. 1c 4^th^ panel). 5) Six double bonds of the chromophore are refined to either *cis* or *trans* configurations in the ground state and all intermediates with small deviations from the ideal geometry. Therefore, each double bond can be fitted with a plane. The fitted plane of each double bond tilts from the ground state as the conformation evolves through the intermediates (Fig. 1c bottom panel). These conformational parameters provide a good metric to evaluate and compare the intermediate species.

In contrast to the flat all-*trans* retinal chromophore in the ground state (Fig. 1c 2^nd^ panel), the side chain of Lys216 is highly twisted forming two near-90° single bonds (Fig. 1c 4^th^ panel), which results in a corner at C_ε_ that deviates dramatically from the plane of the all-*trans* retinal (Fig. 1c 2^nd^ panel). This corner and its interaction with helix C is important to the function of this proton pump (Ren, 2021), which is a topic outside the scope of isomerization sampling here. The refined geometry of the retinal in I’ retains a near perfect all-*trans* configuration, including the Schiff base (SB) double bond C_15_=N_ζ_, while various single bonds along the polyene chain deviate from the standard *anti* conformation significantly (Fig. 1c 4^th^ panel). The torsional deviations from *anti* are in a descending order from the **β**-ionone ring to the SB. However, the largest torsional twist to 90° is found at the single bond C_δ_-C_γ_ in the Lys216 anchor (Fig. 1c 4^th^ panel). These torsional changes at single bonds are among the earliest conformational responses to photon absorption due to the low cost in energy. On the contrary, no immediate change is visible at the double bonds (see below). The torsional changes of the single bonds result in an S-shaped retinal shortened by ~4% (Fig. 1c 3^rd^ panel). The distal segment C_6_-C_12_ moves inboard up to 0.9 Å and the proximal segment C_13_-C_ε_, including the SB, moves outboard up to 1.6 Å (Fig. 1c 1^st^ and 2^nd^ panels). This creased retinal observed here at around 30 fs (Fig. 2d) is attributed to the direct consequence of a compression under an attraction force between the β-ionone ring and the SB (see below).

The refined structure of the I state (Fig. S7) shows that the retinal remains in near perfect all-*trans*, including the SB, and as creased as its precursor I’ (Fig. 1b). The torsional deviations from *anti* single bonds become even more severe compared to the I’ state and remain in a descending order from the β-ionone ring to the SB (Fig. 1c 4^th^ panel). The major difference from its precursor is that the single bond N_ζ_-C_ε_ now adopts a perfect *syn* conformation, and C_δ_-C_γ_ is restored to perfect *anti* (Figs. 1c 4^th^ panel and 3c). As a result, the anchor Lys216 has largely returned to its resting conformation. This prompt restoration of the resting conformation is found important to the pumping mechanism of bR presented in the companion paper (Ren, 2021). Such a lack of substantial change between the ground state and the intermediate I was previously noted by a comparison of the native retinal to a chemically locked C_13_=C_14_ double bond (Zhong et al., 1996).

Remarkably, the major component ***U***_10_ reconstituted into the difference map of I – bR contains widespread signal associated with all seven helices (Fig. 3c). The reconstituted map clearly shows collective outward motions from the center (Fig. 3d) suggesting an expansion of the retinal binding pocket at hundreds of fs, which is confirmed by the refined structure of the I state (Fig. 3e top panel). For example, the distances between the C_α_ atoms increase by 0.8 Å between Arg82 and Phe208 and by 0.7 Å between Tyr83 and Trp182. It is noteworthy that similar signals in the protein are present in the raw difference map calculated from the time point of 457-646 fs from Nogly et al. (6g7j) prior to an SVD analysis (Fig. S8). The refined structure also shows the expansion to a lesser extent (Fig. S9 top panel). But this expansion of the retinal binding pocket was not reported (Nogly et al., 2018).

Transient bleaching at near UV of 265-280 nm was observed before 200 fs and attributed to structural changes in the retinal skeleton and the surrounding Trp residues (Schenkl et al., 2005). Recent deep-UV stimulated Raman spectroscopy also demonstrated that motions of Trp and Tyr residues start to emerge at 200 fs and remain steady until the isomerization is over at 30 ps (Tahara et al., 2019). Here the refined structure of the I state with displaced helices and an expanded retinal binding pocket offers an explanation for the stimulated Raman gain change at hundreds of fs. However, it is unclear why and how such extensive protein responses take place even before the retinal isomerization. According to the broadly accepted concept of proteinquake, initial motions are generated at the epicenter where the chromophore absorbs a photon and then propagated throughout the protein matrix (Ansari et al., 1985). It is plausible that these ultrafast protein responses are the direct consequence of isomerization sampling in a confined protein pocket. It was observed in organic solvents using high-pressure liquid chromatography (HPLC) that all-*trans* retinal could isomerize at various double bonds along the polyene chain to adopt 9-, 11-, and 13-*cis* configurations, but with rather poor quantum yields (Freedman and Becker, 1986; Koyama et al., 1991). This intrinsic property of the all-*trans* retinal would behave the same even when it is incorporated in the protein except that the protein matrix herds the chromophores on the right track of the productive photocycle and keeps the concentrations of the attempted byproducts low. These byproduct conformations of the retinal during isomerization sampling are too numerous and too minor to be observed experimentally. Nevertheless, they cause a common effect, an expansion of its binding pocket, since the all-*trans* retinal in the resting state is tightly boxed by massive side chains all around (Figs. 3a and S10). Any attempt to isomerize would push against this box one way or another. For instance, triple attempts to isomerize simultaneously at 11, 13, and 15 positions were suggested by a quantum mechanics/molecular mechanics simulation (Altoè et al., 2010). A molecular dynamics simulation also concluded that the protein arrests many different photoisomerization paths and funnels them into a single, productive path (Hayashi et al., 2003). When the retinal binding pocket is altered in mutants, the quantum yield of each isomerization byproduct is expected to increase resulting in an impaired productive pathway (see below).

### Intermediates J’, J and productive isomerization of retinal

The time point of 10 ps of Nogly et al. (6g7k) differs from the previous time point of 457-646 fs (6g7j) by negating the component of ***U***_10_ (Fig. 1a), which leads to a restoration of the normal retinal binding pocket in J’ from an expanded one in the I state followed by a contraction in J (Fig. 3e bottom panel). Two time-independent structures of J’ (green; Fig. S11) and J (gray; Fig. S12) are refined with the same protocol based on the respective reconstituted difference maps J’ – bR = 2700***U***_14_ - 1300***U***_17_ and J – bR = -4200***U***_10_ + 2000***U***_14_ - 300***U***_17_ (Methods). Their populations peak at the approximate time of ~700 fs and ~20 ps, respectively. The observed contraction of the retinal binding pocket is likely due to an elastic recoiling of the seven helical bundle following its transient expansion caused by the isomerization sampling. A slight contraction is also present in the refined structure of 6g7k at 10 ps (Fig. S9 bottom panel).

The creased retinal persists in J’ but starts to relax back to a flat conformation in J (Fig. 1b and 1c 2^nd^ panel). The difference map of J’ – bR clearly shows the 13-*cis* configuration (Fig. 4a). Indeed, near perfect 13-*cis* is successfully refined in both J’ and J structures (Fig. 1c 4^th^ panel). While the SB double bond C_15_=N_ζ_ is momentarily distorted from the *trans* configuration in J’ with a torsion angle of 133°, a perfect *trans* configuration at C_1_5=N_ζ_ is promptly restored in J (Fig. 1c 4^th^ panel). The refined structures of this series of early intermediates show that the SB N_ζ_ is rotating clockwise in the entire process of the isomerization of I’ → I → J’ → J, if the retinal is viewed from the proximal to distal direction. It seems that the isomerization starts in an expanded retinal binding pocket and finishes in a tighter one. Whether the pocket expansion and contraction are required for the productive isomerization and what role the low frequency oscillations play in isomerization will need more time points at short delays to further isolate the molecular events temporally.

**Figure 4.**
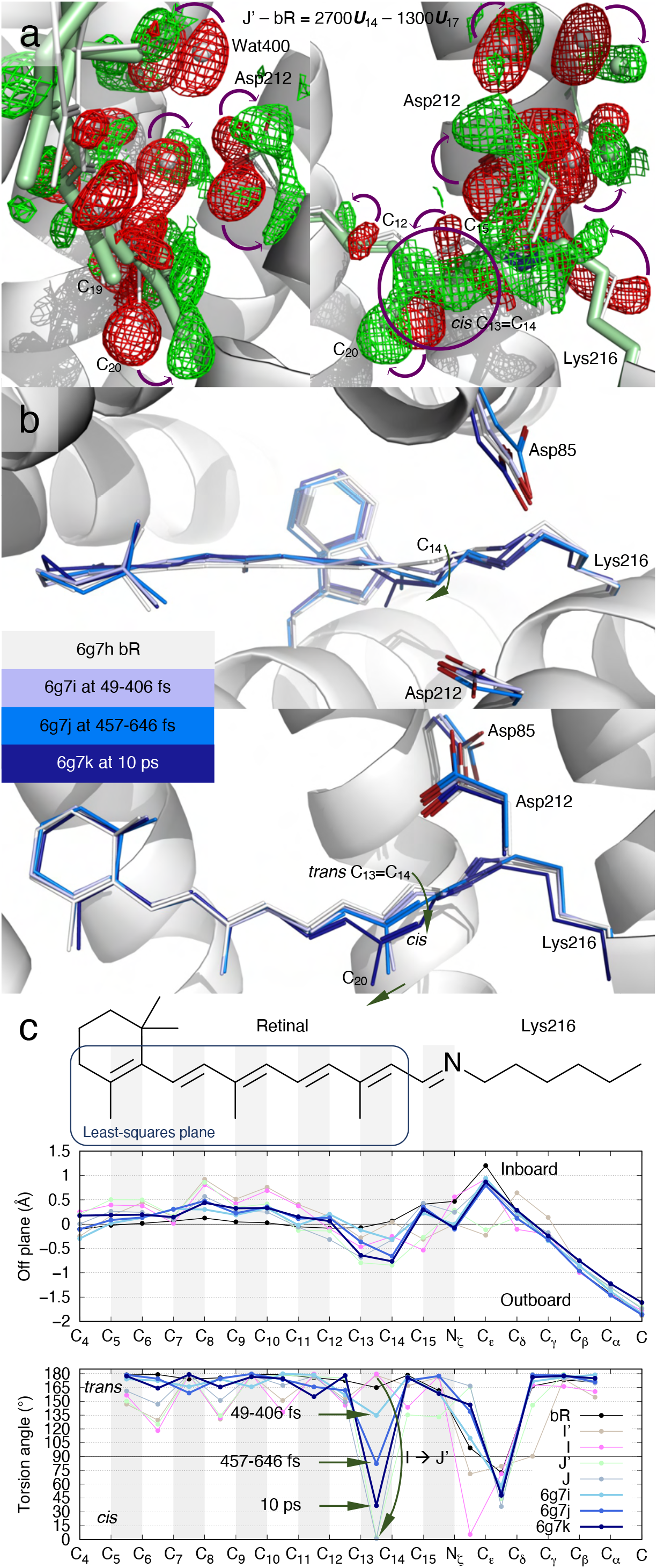
Photoisomerization at C_13_=C_14_ of retinal in bR. (a) Reconstituted difference map J’ – bR from ***U***_14_ and ***U***_17_ (Figs. 2ab and S5). The map is contoured at ±3.5*σ* in green and red mesh, respectively. These difference densities are the direct evidence of isomerization at hundreds of fs. The refined structure of J’ in 13-*cis* is in green. C_13_=C_14_ in perfect *cis* is shown in the circled green positive density. J’ is the earliest 13-*cis* species at hundreds of fs. (b) Published structures of Nogly et al. The chromophore conformations in darker and darker blues are overlayed with the resting state in white. Photoisomerization is interpreted as a gradual change in conformation from tens to hundreds of fs to ps. Even at 10 ps, C_13_=C_14_ is not yet in perfect *cis* (Nogly et al., 2018). (c) Conformational parameters calculated from the published structures of Nogly et al. The chemical structure of the chromophore on top is aligned to the horizontal axis. Double bonds are shaded in gray. 1) A plane is least-squares fitted to C_4_ through C_14_ of the resting state as boxed on the chemical structure. The distances of all atoms to this plane in the inboard and outboard directions show slight crease of the chromophore captured by the structures of Nogly et al. (top panel). 2) The torsion angles indicate no significant rotation of single bonds captured by the structures of Nogly et al. (bottom panel). However, C_13_=C_14_ is transitioning gradually from *trans* to *cis*. In contrast, the refined structures of I and J’ show a complete photoisomerization from perfect *trans* to perfect *cis* directly.

The previous interpretations of the mixed signals of multiple intermediate species presented a rotation of the C_14_ atom, instead of N_ζ_, in the counterclockwise direction if viewed from the proximal end of the retinal (Fig. 4b) (Kovacs et al., 2019; Nogly et al., 2018). This rotation starts at the time point of 49-406 fs (6g7i) and proceeds continuously through hundreds of fs to ps. At 10 ps time point (6g7k), this rotation still does not complete and results in the remaining twist of 37° in the torsion angle of C_13_=C_14_ (Fig. 4c bottom panel). An experimentally determined double bond with a torsion angle of 82°, for example 6g7j, indicates that this twisted double bond conformation is more populated than any other conformation at hundreds of fs. Such interpretation is in direct contradiction of the current view in molecular dynamics. In theory, a double bond conformation spends most time in either energy well of *trans* or *cis*. The transition time between these energy wells is relatively short. A twisted double bond conformation cannot be significantly populated compared to the discrete populations in *trans* and *cis*, that is, cannot be observed, unless an additional energy well emerges near the torsion angle of 90°, which no evidence supports. The gradual rotation of C_13_=C_14_ double bond presented in the refined structures of Nogly et al. and Kovacs et al. was a misinterpretation of the mixed signals of multiple intermediate species as a single conformer. The gradual rotation does not occur in isomerization, rather it merely reflects the gradual population shift from discrete states of *trans* to *cis*. In stark contrast, this work presents that perfect *trans* to perfect *cis* isomerization occurs in a single move during the transition of I → J’ (Fig. 4c bottom panel). Are these two different interpretations of the same data equivalent and inconsequential to mechanistic understanding? The discrete states in *trans* and *cis* presented here result in two sharp U-turns of the SB N_ζ_ not observed in the previous interpretations, the first one at I’ around 30 fs and the second one during I → J’ isomerization at ~500 fs (Fig. 1b inset). The functional implication of these ultrafast SB U-turns is presented in the companion paper (Ren, 2021).

### Coulombic attraction as driving force of isomerization sampling

The fundamental questions remain: What is the driving force that causes the all-*trans* retinal to isomerize after a photon absorption, at several double bonds with a preference at C_11_=C_12_ if not restrained but exclusively at C_13_=C_14_ in bR? Which structural feature(s) of the chromophore and/or its protein environment differentiate the excited state from the ground state? How does the protein environment enhance the quantum yield and speed up the reaction rate of the isomerization to 13-*cis* by tuning the shape of the potential energy surfaces? When does the crossing of the conical intersection occur along the photocycle trajectory? Here I hypothesize that a Coulombic attraction between the β-ionone ring and the SB at the Frank-Condon point, 0+ time point, provides the initial driving force upon a photon absorption. The electric field spectral measurements (Mathies and Stryer, 1976) and the quantum mechanics simulation (Nogly et al., 2018) suggested that a charge separation occurs along the polyene chain at the excited state of bR. Such a dipole moment was also detected through a transient bleaching signal at near UV region (Schenkl et al., 2005). It can be shown that a plausible charge separation of ±0.1e between the β-ionone ring and the SB would cause an attraction force > 1 pN. If calibrated with the measured range of dipole moment of 10-16 D (Mathies and Stryer, 1976), the charge separation could reach the level of ±0.16e to ±0.26e, giving rise to an attraction force of 3.5-9 pN between the β-ionone ring and the SB. This attraction force is evidently sufficient to crease the flat all-*trans* retinal into an S-shape and to compress it slightly within tens of fs as observed in I’ and I states (Figs. 1b, 1c 2^nd^ and 3^rd^ panels, and 2d). In the meanwhile, this very attraction force also triggers simultaneous attempts of double bond isomerizations and single bond rotations along the polyene chain that cause the expansion of the retinal binding pocket as observed at hundreds of fs. Despite such observable expansion, the elastic strength of the seven helical bundle blocks all reaction pathways of the isomerization sampling except the productive one. Following the only successful isomerization at C_13_=C_14_, the chromophore segment from C_15_ to C_δ_ is attracted to the β-ionone ring; and these two parts become significantly closer (Fig. 1c 3^rd^ panel). None of the single bond rotations can complete under the restraints of the protein (Figs. 3a and S10). Especially, the segment closer to the midpoint of the retinal is more confined due to the steric hinderance of Thr90 and Tyr185 from the inboard and outboard sides, respectively (Fig. 3a). Therefore, the single bonds deviate from *anti* less and less towards the midpoint (Fig. 1c 4^th^ panel).

I could further speculate that the charge separation along the polyene chain differentiates the excited state from the ground state. Therefore, the disappearance of any structural effects caused by the charge separation marks the crossing of the conical intersection. The light-induced charge separation seems to subside gradually as the reaction proceeds beyond the K state as indicated by the slow restoration of the perfect *anti* conformation and the reluctant flattening of the retinal plane (Fig. 1c 2^nd^ and 4^th^ panel), where the refined structures of K and L are taken from the companion paper (Ren, 2021). If it is safe to assume that the single bond distortion away from a perfect *anti* and the extent of the crease in the retinal plane are direct measures of the charge separation with a delay no longer than ns time scale, the observed crossing of the conical intersection is surprisingly late during the K → L transition at a few μs. The structural basis of tuning potential energy surface arises from the stereochemical hinderance of the retinal binding pocket that makes all isomerization attempts and single bond rotations energetically costly except one.

Apparently, the same charge separation and the attraction force upon photon absorption also take place in a solution sample of free retinal. Compared to the retinal embedded in protein, photoisomerization in solution is nonspecific, resulting in a range of byproducts, since an isomerization at any position would bring the SB significantly closer to the β-ionone ring. The major photoproduct is 11-*cis* among several possibilities in solution simply because C_11_=C_12_ double bond is located at the midpoint from the β- ionone ring to the SB thus most vulnerable to a mechanical twist. It is also understandable that each of the byproducts could only achieve a poor quantum yield (Freedman and Becker, 1986; Koyama et al., 1991) as rotations at multiple single bonds driven by the same attraction force and achieving a similar kink in the polyene chain would further sidetrack the double bond isomerizations thus diminishing their quantum yields. Since the total quantum yield of isomerization at various double bonds is ~1/4, it can be predicted that the total quantum yield of single bond rotations is ~3/4 in solution. Because the deeper valleys on the potential energy surface leading to single bonds in *syn* are more energetically favorable routes than the shallower valleys leading to *cis* double bonds. However, these byproducts due to single bond rotations are short-lived beyond detection by HPLC as they spontaneously revert back in solution. The protein environment in bR plays a major role in enhancing the quantum yield and accelerating the reaction rate of the isomerization to 13-*cis* by shutting down all other reaction pathways triggered by the same charge separation. This is further elucidated by the mutant functions below.

### Isomerization byproducts permitted by mutant protein environments

The structure of a double mutant T90A/D115A (3cod) showed little difference from the wildtype (Joh et al., 2008) while the single mutants T90V and T90A retain < 70% and < 20% of the proton pumping activity, respectively (Marti et al., 1991; Perálvarez et al., 2001). These observations illustrate that some nonproductive pathways of the isomerization sampling succeed more in the altered retinal binding pocket. In the wildtype structure, Thr90 in helix C points towards the C_11_=C_12_-C_13_-C_20_ segment of the retinal from the inboard with its C_γ_ atom 3.7 Å from the retinal plane (Fig. 3a). Given the van der Waals radius rc of 1.7 Å, only 0.3 Å is spared for the H atoms of the C_γ_ methyl group thereby effectively shutting down the nonproductive pathways of the isomerization sampling. In addition, the C_ε_ methyl group of Met118 and the C_δ_ ethyl group of Ile119 in helix D are 3.7-4.6 Å away from the retinal segment of C_7_=C_8_-C_9_=C_10_ and C_19_ from the inboard (Fig. S10). Any motion of the retinal would have to push helices C and D toward inboard causing an expansion of its binding pocket. Missing the close contact in T90A increases the room to 1.9 Å for isomerization byproducts, which would greatly reduce the quantum yield of the 13-*cis* productive isomerization thus retain < 20% of the activity.

In addition to 13-*cis*, the retinal in the light adapted T90V mutant showed 9- and 11- *cis* configurations at the occupancies of 3% and 18%, respectively, while these configurations were not detected in light adapted wildtype (Marti et al., 1991). Then why would a Val residue at this position with an equivalent C_γ_ methyl group permit the formation of some isomerization byproducts? In wildtype bR, the side chain of Thr90 engages two strong H-bonds Trp86O-Thr90O_γ_–D115O_δ_ so that its C_γ_ methyl group is aligned toward the retinal (Fig. 3b). Without these H-bonds in T90V, the isopropyl group of Val90 is free to adopt other rotameric positions so that neither of the C_γ_ methyl groups has to point directly to the retinal, which increases the available room for the formation of some isomerization byproducts. The conformational change caused by the Val substitution at this position was detected by spectroscopy (Kandori et al., 1999). Compared to the light adapted state with 3% *9-cis* and 18% 11-cis, these isomerization byproducts could reach even higher percentages during active photocycles thus reduce the proton pumping activity below 70%.

It is important that the C_γ_ methyl group of Thr90 in the wildtype is pointing to C_11_=C_12_ double bond located at the midpoint of the retinal as 11-_cis_ is the most probable byproduct (Hamm et al., 1996; Logunov et al., 1996). The potential energy surface of the excited state in solution must feature a major branch of energetic valley leading to 11-*cis*. But this valley is blocked by the stereo hinderance of Thr90 braced by the H-bonds.

From the outboard, the side chain of Tyr185 in helix F is nearly parallel to the mid segment of the retinal plane with a distance of 3.5 Å (Fig. 3a), and Pro186 is also parallel to the distal segment with 3.7-4.4 Å distance (Fig. S10). These close contacts of flat areas from C_5_ to C_14_ of the retinal prevents any significant motion of the retinal toward the outboard. Even slight motions would push helix F away as observed here in the expansion of the retinal binding pocket. The mutant Y185F largely retains the flat contact so that its proton pumping activity does not reduce much (Hackett et al., 1987; Mogi et al., 1987). However, it is predictable that various single mutants at this position with smaller and smaller side chains would promote more and more isomerization byproducts and eventually shut down proton pumping.

Two massive side chains of Trp86 and 182 from the extracellular and cytoplasmic sides respectively do not seem to play a significant role in suppressing byproduct formation as shown by the mutant W182F that retains the most of the wildtype activity (Hackett et al., 1987), since the motions involved in isomerization sampling are oriented more laterally. The transient expansion and contraction of the retinal binding pocket (Fig. 3e) indicate that the tight box surrounds the mid-segment of the retinal (Fig. 3a) is not completely rigid. Rather, its plasticity must carry sufficient strength to prevent isomerization byproducts. Presumably, this strength originates from the mechanical property of the helical bundle.

## Concluding remarks

In summary, this work employs a numerical resolution from concurrent structural events, hence substantially improves the quality of electron density maps of unscrambled intermediate species that are otherwise difficult to observe. This approach reveals the transient structural responses to many unsuccessful attempts of double bond isomerization and single bond rotation. These findings underscore an important implication, that is, a nonspecific Coulombic attraction provides the same driving force for the isomerization sampling with and without a protein matrix. A productive isomerization at a specific double bond is guided by the incorporation of the chromophore in a specific protein environment. The productive pathway is selected from numerous possibilities via stereochemical hinderance. This case study offers a detailed example elucidating how a protein structure achieves regioselectivity in a biochemical reaction that catalyzes a specific double bond isomerization by accelerating its reaction rate and shutting down all other byproduct pathways. Nevertheless, this nonspecific Coulombic attraction force may not be directly applicable to the photoisomerization of retinal from 11-*cis* to all-*trans* in the activation of visual rhodopsins. The commonality among microbial and animal rhodopsins perhaps exists only at a high level in a rather abstract sense, if any. These photoreactions to and from the all-*trans* retinal not necessarily share any common trajectory on the potential energy surfaces. The key difference is bR as an energy convertor versus a visual rhodopsin as a quantum detector (Lewis, 1978).

## Acknowledgements

This work is supported in part by the grant R01EY024363 from National Institutes of Health. The following database and software are used in this work: CCP4 (ccp4.ac.uk), Coot (www2.mrc-lmb.cam.ac.uk/Personal/pemsley/coot), dynamiX™ (Renz Research, Inc.), gnuplot (gnuplot.info), PDB (rcsb.org), PHENIX (phenix-online.org), PyMOL (pymol.org), Python (python.org), and SciPy (scipy.org).

## Data Availability

Depositions of the refined structures of I’, I, J’, and J to PDB-Dev (pdb-dev.wwpdb.org) are underway.

## Methods

From the outset, the key presumption is that every crystallographic dataset, at a given temperature and a given time delay after the triggering of a photochemical reaction, captures a mixture of unknown number of intermediate species at unknown fractions. Needless to say, all structures of the intermediates are also unknown except the structure at the ground state that has been determined and well refined by static crystallography. A simultaneous solution of all these unknowns requires multiple datasets that are collected at various temperatures or time delays so that a common set of intermediate structures are present in these datasets with variable ratios. If the number of available datasets is far greater than the number of unknowns, a linear system can be established to overdetermine the unknowns with the necessary stereochemical restraints (Ren et al., 2013). The analytical methods used in this work to achieve such overdetermination have been incrementally developed in the past years and recently applied to another joint analysis of the datasets of carbonmonoxy myoglobin (Ren, 2019). Time-resolved datasets collected with ultrashort pulses from an X-ray free electron laser were successfully analyzed by these methods to visualize electron density components that reveal transient heating, 3*d* electrons of the heme iron, and global vibrational motions. This analytical strategy is recapped below.

The methodological advance in this work is the refinement of each pure intermediate structure that has been deconvoluted from multiple mixtures. Structure factor amplitudes of a single conformation free of heterogeneity are overdetermined. Given the deconvoluted structure factor amplitude set of a pure state, the standard structural refinement software with the built-in stereochemical constraints is taken full advantage of, e.g. PHENIX (Adams et al., 2010; Liebschner et al., 2019). In case that the computed deconvolution has not achieved a single pure structural species, the structural refinement is expected to make such indication.

### Difference Fourier maps

A difference Fourier map is synthesized from a Fourier coefficient set of *F*_light_-*F*_reference_ with the best available phase set, often from the ground state structure. Before Fourier synthesis, *F*_light_ and *F*_reference_ must be properly scaled to the same level so that the distribution of difference values is centered at zero and not skewed either way. A weighting scheme proven effective assumes that a greater amplitude of a difference Fourier coefficient *F*_light_-*F*_reference_ is more likely caused by noise than by signal (Ren et al., 2001, 2013; Šrajer et al., 2001; Ursby and Bourgeois, 1997). Both the dark and light datasets can serve as a reference in difference maps. If a light dataset at a certain delay is chosen as a reference, the difference map shows the changes since that delay time but not the changes prior to that delay. However, both the dark and light datasets must be collected in the same experiment. A cross reference from a different experimental setting usually causes large systematic errors in the difference map that would swamp the desired signals. Each difference map is masked 3.5 Å around the entire molecule of bacteriorhodopsin (bR). No lipid density is analyzed.

### Meta-analysis of protein structures

Structural meta-analysis based on singular value decomposition (SVD) has been conducted in two forms. In one of them, an interatomic distance matrix is calculated from each protein structure in a related collection. SVD of a data matrix consists of these distance matrices enables a large-scale joint structural comparison but requires no structural alignment (Ren, 2013a, 2013b, 2016). In the second form, SVD is performed on a data matrix of electron densities of related protein structures (Ren, 2019; Ren et al., 2013; Schmidt et al., 2003, 2010). Both difference electron density maps that require a reference dataset from an isomorphous crystal form and simulated annealing omit maps that do not require the same unit cell and space group of the crystals are possible choices in a structural meta-analysis (Ren, 2019; Ren et al., 2013). The interatomic distances or the electron densities that SVD is performed on are called core data. Each distance matrix or electron density map is associated with some metadata that describe the experimental conditions under which the core data are obtained, such as temperature, pH, light illumination, time delay, mutation, etc. These metadata do not enter the SVD procedure. However, they play important role in the subsequent interpretation of the SVD result. This computational method of structural analysis takes advantage of a mathematical, yet practical, definition of conformational space with limited dimensionality (Ren, 2013a). Each experimentally determined structure is a snapshot of the protein structure. A large number of such snapshots taken under a variety of experimental conditions, the metadata, would collectively provide a survey of the accessible conformational space of the protein structure and reveal its rection trajectory. Such joint analytical strategy would not be effective in early years when far fewer protein structures were determined to atomic resolution. Recent rapid growth in protein crystallography, such as in structural genomics (Berman et al., 2012; Bonvin, 2021; Chandonia and Brenner, 2006) and in serial crystallography (Glynn and Rodriguez, 2019; Schaffer et al., 2021), has supplied the necessarily wide sampling of protein structures for a joint analytical strategy to come of age. The vacancies or gaps in a conformational space between well-populated conformational clusters often correspond to less stable transient states whose conformations are difficult to capture, if not impossible. These conformations are often key to mechanistic understanding and could be explored by a back calculation based on molecular distance geometry (Ren, 2013a, 2016), the chief computational algorithm in nucleic magnetic resonance spectroscopy (NMR), and by a structure refinement based on reconstituted dataset, a major methodological advance in this work (see below). These structures refined to atomic resolution against reconstituted datasets may reveal short-lived intermediate conformation hard to be captured experimentally. Unfortunately, a protein structure refined against a reconstituted dataset currently cannot be recognized by the Protein Data Bank (PDB). Because crystallographic refinement of a macromolecular structure is narrowly defined as a correspondence from one dataset to one structure. A never-observed dataset reconstituted from a collection of experimental datasets does not match the well-established crystallographic template of PDB; let alone a refinement of crystal structure with the NMR algorithm.

A distance matrix contains *M* pairwise interatomic distances of a structure in the form of Cartesian coordinates of all observed atoms. An everyday example of distance matrix is an intercity mileage chart appended to the road atlas. Differences in the molecular orientation, choice of origin, and crystal lattice among all experimentally determined structures have no contribution to the distance matrices. Due to its symmetry, only the lower triangle is necessary. A far more intimate examination of protein structures in PDB is a direct analysis of their electron density maps instead of the atomic coordinates. *M* such (difference) electron densities, often called voxels in computer graphics, are selected by a mask of interest. In the case of difference maps, only the best refined protein structure in the entire collection supplies a phase set for Fourier synthesis of electron density maps. This best structure is often the ground state structure determined by static crystallography. Other refined atomic coordinates from the PDB entries are not considered in the meta-analysis. That is to say, a meta-analysis of difference electron density maps starts from the X-ray diffraction data archived in PDB rather than the atomic coordinates interpreted from the diffraction data, which removes any potential model bias.

### Singular value decomposition of (difference) electron density maps

An electron density map, particularly a difference map as emphasized here, consists of density values on an array of grid points within a mask of interest. All *M* grid points in a three-dimensional map can be serialized into a one-dimensional sequence of density values according to a specific protocol. It is not important what the protocol is as long as a consistent protocol is used to serialize all maps of the same grid setting and size, and a reverse protocol is available to erect a three-dimensional map from a sequence of *M* densities. Therefore, a set of *N* serialized maps, also known as vectors in linear algebra, can fill the columns of a data matrix **A** with no specific order, so that the width of **A** is *N* columns, and the length is *M* rows. Often, *M* >> *N*, thus **A** is an elongated matrix. If a consistent protocol of serialization is used, the corresponding voxel in all *N* maps occupies a single row of matrix **A**. This strict correspondence in a row of matrix **A** is important. Changes of the density values in a row from one structure to another are due to either signals, systematic errors, or noises. Although the order of columns in matrix **A** is unimportant, needless to say, the metadata associated with each column must remain in good bookkeeping.

SVD of the data matrix **A** results in **A** = **UWV**^T^, also known as matrix factorization. Matrix **U** has the same shape as **A**, that is, *N* columns and *M* rows. The *N* columns contain decomposed basis components ***U***_*k*_, known as left singular vectors of *M* items, where *k* = 1, 2,…, *N*. Therefore, each component ***U***_*k*_ can be erected using the reverse protocol to form a three-dimensional map. This decomposed elemental map can be presented in the same way as the original maps, for example, rendered in molecular graphics software such as Coot and PyMol. It is worth noting that these decomposed elemental maps or map components ***U***_*k*_ are independent of any metadata. That is to say, these components remain constant when the metadata vary. Since each left singular vector ***U***_*k*_ has a unit length due to the orthonormal property of SVD (see below), that is, |***U***_*k*_| = 1, the root mean squares (rms) of the items in a left singular vector is 1/√*M* that measures the quadratic mean of the items.

The second matrix **W** is a square matrix that contains all zeros except for *N* positive values on its major diagonal, known as singular values *w_k_*. The magnitude of *w_k_* is considered as a weight or significance of its corresponding component ***U***_*k*_. The third matrix **V** is also a square matrix of *N* × *N*. Each column of **V** or row of its transpose **V**^T^, known as a right singular vector ***V***_*k*_, contains the relative compositions of ***U***_*k*_ in each of the *N* original maps. Therefore, each right singular vector **V**_*k*_ can be considered as a function of the metadata. Right singular vectors also have the same unit length, that is, |**V**_*k*_| = 1. Effectively, SVD separates the constant components independent of the metadata from the compositions that depend on the metadata.

A singular triplet denotes 1) a decomposed component ***U***_*k*_, 2) its singular value *w_k_*, and 3) the composition function **V**_*k*_. Singular triplets are often sorted in a descending order of their singular values *w_k_*. Only a small number of *n* significant singular triplets identified by the greatest singular values *w*_1_ through *w_n_* can be used in a linear combination to reconstitute a set of composite maps that closely resemble the original ones in matrix **A**, where *n* < *N*. For example, the original map in the ith column of matrix **A** under a certain experimental condition can be closely represented by the ith composite map *w*_1_*v*_1*i*_***U***_1_ + *w*_2_*v*_2*i*_***U***_2_ + … + *w_n_v_ni_****U***_*n*_, where (*v*_1*i*_, *v*_2*i*_,…) is from the *i*th row of matrix **V**. The coefficient set for the linear combination is redefined here as *c_ki_* = *w_k_v_ki_*/√*M*. The rms of the values in a map component, or the average magnitude measured by the quadratic mean, acts as a constant scale factor that resets the modified coefficients *c_ki_* back to the original scale of the core data, such as Å for distance matrices and e-/Å^3^ for electron density maps if these units are used in the original matrix **A**. Practically, an electron density value usually carries an arbitrary unit without a calibration, which makes this scale factor unnecessary. In the linear combination *c*_1*i*_***U***_1_ + *c*_2*i*_***U***_2_ + … + *c_ni_****U***_*n*_, each component ***U***_*k*_ is independent of the metadata while how much of each component is required for the approximation, that is, *c_ki_*, depends on the metadata.

Excluding the components after ***U***_*n*_ in this approximation is based on an assumption that the singular values after *w_n_* are very small relative to those from *w*_1_ through *w_n_*. As a result, the structural information evenly distributed in all *N* original maps is effectively concentrated into a far fewer number of *n* significant components, known as information concentration or dimension reduction. On the other hand, the trailing components in matrix **U** contain inconsistent fluctuations and random noises. Excluding these components effectively rejects noises (Schmidt et al., 2003). The least-squares property of SVD guarantees that the rejected trailing components sums up to the least squares of the discrepancies between the original core data and the approximation using the accepted components.

However, no clear boundary is guaranteed between signals, systematic errors, and noises. Systematic errors could be more significant than the desired signals. Therefore, excluding some components from 1 through *n* is also possible. If systematic errors are correctly identified, the reconstituted map without these significant components would no longer carry the systematic errors.

### The orthonormal property of SVD

The solution set of SVD must guarantee that the columns in **U** and **V**, the left and right singular vectors ***U***_*k*_ and ***V***_*k*_, are orthonormal, that is, ***U***_*h*_∂***U***_*k*_ = ***V***_*h*_∂***V***_*k*_ = 0 (ortho) and ***U***_*k*_∂***U***_*k*_ = ***V***_*k*_∂***V***_*k*_ = 1 (normal), where *h* ≠ *k* but both are from 1 to *N*. The orthonormal property also holds for the row vectors. As a result, each component ***U***_*k*_ is independent of the other components. In other words, a component cannot be represented by a linear combination of any other components. However, two physical or chemical parameters in the metadata, such as temperature and pH, may cause different changes to a structure. These changes are not necessarily orthogonal. They could exhibit some correlation. Therefore, the decomposed components ***U***_*k*_ not necessarily represent any physically or chemically meaningful changes (see below).

Due to the orthonormal property of SVD, an *N*-dimensional Euclidean space is established, and the first *n* dimensions define its most significant subspace. Each coefficient set ***c***_*i*_ = (*c*_1*i*_, *c*_2*i*_,…,*c_ni_*) of the ith composite map is located in this *n*-dimensional subspace. All coefficient sets for *i* = 1, 2, …, *N* in different linear combinations to approximate the *N* original maps in a least-squares sense can be represented by *N* points or vectors ***c***_1_, ***c***_2_, …, ***c***_*N*_ in the Euclidean subspace. This n-dimensional subspace is essentially the conformational space as surveyed by the jointly analyzed core data. The conformational space is presented as scatter plots with each captured structure represented as a dot located at a position determined by the coefficient set **c***i* of the *i*th observed map. When the subspace has greater dimensionality than two, multiple two-dimensional orthographical projections of the subspace are presented, such as Fig. 1a. These scatter plots are highly informative to reveal the relationship between the (difference) electron density maps and their metadata.

If two coefficient sets ***c***_*i*_ ≈ ***c***_*j*_, they are located close to each other in the conformational space. Therefore, these two structures *i* and *j* share two similar conformations. Two structures located far apart from each other in the conformational space are dissimilar in their conformations, and distinct in the compositions of the map components. A reaction trajectory emerges in this conformational space if the temporal order of the core data is experimentally determined (Fig. 1a). Otherwise, an order could be assigned to these structures based on an assumed smoothness of conformational changes along a reaction trajectory (Ren, 2013a, 2013b, 2016). Causation and consequence of structural motions could be revealed from the order of the structures in a series, which may further lead to structural mechanism. In addition, an off-trajectory location in the conformational space or a location between two clusters of observed structures represents a structure in a unique conformation that has never been experimentally captured. Such a hypothetical structure can be refined against a reconstituted distance matrix using molecular distance geometry (Ren, 2013a, 2013b, 2016) or a reconstituted electron density map with the method proposed below.

### Rotation in SVD space

Dimension reduction is indeed effective in meta-analysis of protein structures when many datasets are evaluated at the same time. However, the default solution set of SVD carries complicated physical and chemical meanings that are not immediately obvious. The interpretation of a basis component ***U***_*k*_, that is, “what-does-it-mean”, requires a clear demonstration of the relationship between the core data and their metadata. The outcome of SVD does not guarantee any physical meaning in a basis component. Therefore, SVD alone provides no direct answer to “what-does-it-mean”, thus its usefulness is very limited to merely a mathematical construction. However, the factorized set of matrices **U, W**, and **V** from SVD is not a unique solution. That is to say, they are not the only solution to factorize matrix **A**. Therefore, it is very important to find one or more alternative solution sets that are physically meaningful to elucidate a structural interpretation. The concept of a rotation after SVD was introduced by Henry & Hofrichter (Henry and Hofrichter, 1992). But they suggested a protocol that fails to preserve the orthonormal and least-squares properties of SVD. The rotation protocol suggested by Ren incorporates the metadata into the analysis and combines with SVD of the core data. This rotation achieves a numerical deconvolution of multiple physical and chemical factors after a pure mathematical decomposition, and therefore, provides a route to answer the question of “what-does-it-mean” (Ren, 2019). This rotation shall not be confused with a rotation in the three-dimensional real space, in which a molecular structure resides.

A rotation in the *n*-dimensional Euclidean subspace is necessary to change the perspective before a clear relationship emerges to elucidate scientific findings. It is shown below that two linear combinations are identical before and after a rotation applied to both the basis components and their coefficients in a two-dimensional subspace of *h* and *k*. That is,

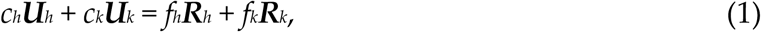

where *c_h_* and *c_k_* are the coefficients of the basis components ***U***_*h*_ and ***U***_*k*_ before the rotation; and *f_h_* and *f_k_* are the coefficients of the rotated basis components ***R***_*h*_ and ***R***_*k*_, respectively. The same Givens rotation of an angle *θ* is applied to both the components and their coefficients:

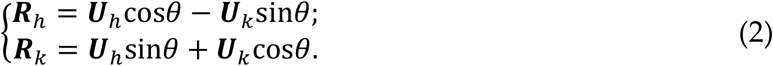

Obviously, the rotated components ***R***_*h*_ and ***R***_*k*_ remain mutually orthonormal and orthonormal to other components. And

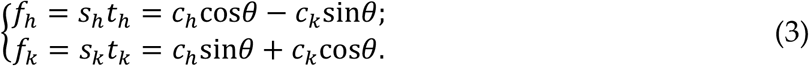

Here 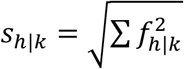 are the singular values that replace *wh* and *w_k_*, respectively, after the rotation. They may increase or decrease compared to the original singular values so that the descending order of the singular values no longer holds. ***T***_*h*|*k*_ = (*t*_*h*|*k*1_, *t*_*h*|*k*2_,…, *t*_*h*|*kN*_) = (*f*_*h*|*k*1_, *f*_*h*|*k*2_,…, *f*_*h*|*kN*_)/*S*_*h*|*k*_ are the right singular vectors that replace ***V***_*h*_ and ***V***_*k*_, respectively. ***T***_*h*_ and ***T***_*k*_ remain mutually orthonormal after the rotation and orthonormal to other right singular vectors that are not involved in the rotation.

Eq. 1 holds because the dot product of two vectors does not change after both vectors rotate the same angle. To prove Eq. 1 in more detail, Eqs. 2 and 3 are combined and expanded. All cross terms of sine and cosine are self-canceled:

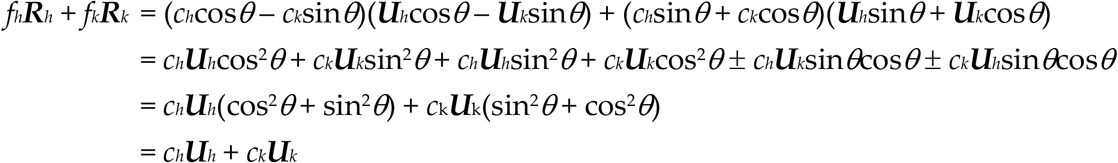

A rotation in two-dimensional subspace of *h* and *k* has no effect in other dimensions, as the orthonormal property of SVD guarantees. Multiple steps of rotations can be carried out in many two-dimensional subspaces consecutively to achieve a multi-dimensional rotation. A new solution set derived from a rotation retains the orthonormal property of SVD. The rotation in the Euclidean subspace established by SVD does not change the comparison among the core data of protein structures. Rather it converts one solution set **A** = **UWV**^T^ to other alternative solutions **A** = **RST**^T^ so that an appropriate perspective can be found to elucidate the relationship between the core data and metadata clearly and concisely.

For example, if one physical parameter could be reoriented along a single dimension *k* but not involving other dimensions by a rotation, it can be convincingly shown that the left singular vector **U***k* of this dimension illustrates the structural impact by this physical parameter. Before this rotation, the same physical parameter may appear to cause structural variations along several dimensions, which leads to a difficult interpretation. Would a proper rotation establish a one-on-one correspondence from all physical or chemical parameters to all the dimensions? It depends on whether each parameter induces an orthogonal structural change, that is, whether structural responses to different parameters are independent or correlated among one another. If structural changes are indeed orthogonal, it should be possible to find a proper rotation to cleanly separate them in different dimensions. Otherwise, two different rotations are necessary to isolate two correlated responses, but one at a time.

For another example, if the observed core datasets form two clusters in the conformational space, a rotation would be desirable to separate these clusters along a single dimension *k* but to align these clusters along other dimensions. Therefore, the component ***U***_*k*_ is clearly due to the structural transition from one cluster to the other. Without a proper rotation, the difference between these clusters could be complicated with multiple dimensions involved. A deterministic solution depends on whether a clear correlation exists between the core data and metadata. A proper rotation may require a user decision. A wrong choice of rotation may select a viewpoint that hinders a concise conclusion. However, it would not alter the shape of the reaction trajectory, nor create or eliminate an intrinsic structural feature. A wrong choice of rotation cannot eliminate the fact that a large gap exists between two clusters of observed core datasets except that these clusters are not obvious from that viewpoint. A different rotation may reorient the perspective along another direction. But the structural conclusion would be equivalent. See example of before and after a rotation in (Ren, 2016).

This rotation procedure finally connects the core crystallographic datasets to the metadata of experimental conditions and accomplishes the deconvolution of physical or chemical factors that are not always orthogonal to one another after a mathematical decomposition. SVD analysis presented in this paper employs rotations extensively except that no distinction is made in the symbols of components and coefficients before and after a rotation except in this section. This method is widely applicable in large-scale structural comparisons. Furthermore, Ren rotation after SVD is not limited to crystallography and may impact other fields wherever SVD is used. For example, SVD is frequently applied to spectroscopic data, images, and genetic sequence data.

### Structural refinement against reconstituted dataset

The linear combination Δ*ρ*(*t*) = *f*_1_(*t*)***R***_1_ + *f*_2_(*t*)***R***_2_ + … + *f_n_*(*t*)***R***_*n*_ after a rotation reconstitutes one of the observed difference maps at a specific time point *t*. This time-dependent difference map depicts an ever-evolving mixture of many excited species. A reconstituted difference map Δ*ρ*(E) for a time-independent, pure, excited species E = intermediate I’, I, J’, and J deconvoluted from many mixtures would take the same form except that only one or very few coefficients remain nonzero if a proper rotation has been found (Table S2). In order to take advantage of the mature refinement software for macromolecular structures with extensive stereochemical restraints, a set of structure factor amplitudes is needed. Therefore, it is necessary to reconstitute a set of structure factor amplitudes that would produce the target difference map Δ*ρ*(E) based on a known structure at the ground state. First, an electron density map of the structure at the ground state is calculated. This calculated map is used as a base map. Second, this base map of the ground state is combined with the positive and negative densities in the target difference map Δ*ρ*(E) so that the electron densities at the ground state are skewed toward the intermediate state. Third, structure factors are calculated from the combined map. Finally, the phase set of the calculated structure factors is discarded, and the amplitudes are used to refine a single conformation of the intermediate species E that Δ*ρ*(E) represents.

This protocol following the SVD and Ren rotation of components achieves a refinement of a pure structural species without the need of alternative conformations. Several points are noteworthy. First, the minimization protocol in this refinement is performed against a numerically reconstituted amplitude set that has never been directly measured from a crystal. This reconstituted dataset could be considered as an extrapolated dataset “on steroids” if compared to the traditional extrapolation of small differences, such as, the Fourier coefficient set to calculate a 3Fo-2Fc map, a technique often used to overcome a partial occupancy of an intermediate structure. An extrapolation of small differences is not directly observed either but computed by an exaggeration of the observed difference based on an assumption that the intermediate state is partially occupied, such as the doubling of the observed difference in 3Fo-2Fc = Fo + 2(Fo-Fc). In contrast to the conventional technique of extrapolation, the deconvolution method applied here is an interpolation among many experimental datasets rather than an extrapolation. Secondly, the deconvolution is a simultaneous solution of multiple intermediate states mixed together instead of solving a single excited state.

Second, a map calculated from the ground state structure is chosen as the base map instead of an experimental map such as Fo or 2Fo-Fc map. If the second step of the protocol is skipped, that is, no difference map is combined with the ground state map, the refinement would result in an *R* factor of nearly zero, since the refinement is essentially against the calculated structure factors (bR in Table S2). This is to say, the residuals of the refinement are solely due to the difference component instead of the base map. This is desirable since errors in the static structure of the ground state are gauged during its own refinement. On the other hand, if an experimental map is chosen as a base map, the refinement *R* factors would reflect errors in both the base map and the difference map, which leads to a difficulty in an objective evaluation of this refinement protocol.

Third, the combination of the base map and a difference map is intended to represent a pure intermediate species. Therefore, alternative conformations in structural refinement that model a mixture of species would defeat this purpose. However, this combined map could be very noisy and may not represent a single species without a proper rotation. This is particular the case, if the target difference map Δ*ρ* is not derived from an SVD analysis and Ren rotation. The SVD analysis identifies many density components that are inconsistent among all observed difference maps and excludes them, which greatly reduces the noise content. Therefore, this refinement protocol may not be very successful without an SVD analysis. Another source of noise originates from the phase set of the structure factors. Prior to the refinement of the intermediate structure, the phase set remains identical to that of the ground state. This is far from the reality when an intermediate structure involves widespread changes, such as those refined in this study. If the rotation after SVD is not properly selected, the target difference map would remain as a mixture minus the ground state. Therefore, the refinement of a single conformation would encounter difficulty or significant residuals, as judged by the *R* factors, the residual map, and the refined structure. A proper solution to this problem is a better SVD solution by Ren rotation rather than alternative conformations. A successful refinement of near perfect *trans* or *cis* double bonds is a good sign to indicate that the reconstituted amplitude set after a rotation reflects a relatively homogeneous structure. If a double bond could not be refined well to near perfect *trans* or *cis* configuration, the dataset of structure factor amplitudes is likely from a mixture of heterogeneous configurations, which occurred frequently in previous studies of bR and photoactive yellow protein (Jung et al., 2013; Lanyi and Schobert, 2007; Nogly et al., 2018). It has been a great difficulty in crystallographic refinement in general that a heterogeneous mixture of conformations cannot be unambiguously refined even with alternative conformations. This difficulty becomes more severe when a mixture involves more than two conformations or when some conformations are very minor.

Lastly, the refinement protocol proposed here could be carried out in the original unit cell and space group of the crystal at the ground state. However, this is not always applicable as the original goal of the meta-analysis is a joint examination of all available structures from a variety of crystal forms. It would be highly desirable to evaluate difference maps of the same or similar proteins from non-isomorphous crystals together by SVD. Alternatively, the refinement protocol could also be performed in the space group of P1 with a virtual unit cell large enough to hold the structure, which is the option in this study (Table S2). This is to say, the entire analysis of SVD-rotation- refinement presented here could be extracted and isolated from the original crystal lattices, which paves the way to future applications to structural data acquired by experimental techniques beyond crystallography, most attractively, to single particle reconstruction in cryo electron microscopy.

## Supplementary Tables

**Table S1.** Datasets analyzed in this work

| Publication | PDB | Label | Resolution | Main conclusions | New findings in this work |
| --- | --- | --- | --- | --- | --- |
| Nogly et al.<br>Science 361,<br>eaat0094,<br>2018 | 6g7h | dark6 | 1.5 Å | Retinal fully<br>isomerizes by 10 ps.<br>But the SB water<br>dissociates earlier. | The short-delay datasets contribute to the<br>structures of $I' \rightarrow I \rightarrow J' \rightarrow J$. Photoisomerization<br>in $J'$; retinal binding pocket expansion before 1 ps<br>in I and contraction at ~20 ps in J |
|  | 6g7i | 49-406fs | 1.9 Å |  |  |
|  | 6g7j | 457-646fs | 1.9 Å |  |  |
|  | 6g7k | 10ps | 1.9 Å |  |  |
| Kovacs et al.<br>Nat.<br>Commun. 10,<br>3177, 2019 | 6ga1 | dark1 | 1.7 Å | The exceedingly high<br>power density of the<br>pump laser causes<br>two-photon<br>absorption.<br>Vibrational motions<br>were observed. | The sub-ps datasets exhibit extensive vibrations<br>at various frequencies. The vibrational signals<br>are widespread over the entire bR molecule and<br>not associated with any structural elements.<br>Therefore, it is concluded that these global<br>vibrations are intrinsic properties of bR induced<br>by short laser pulses. The vibrational signals are<br>more prominent under higher power density of<br>the laser pulses. However, these vibrations are<br>irrelevant to the light-driven proton pumping<br>function of bR. |
|  | 6ga2 | dark2 | 1.8 Å |  |  |
|  | 6rmk | dark3 | 1.8 Å |  |  |
|  | 6ga7 | 240fs | 1.8 Å |  |  |
|  | 6ga8 | 330fs | 1.8 Å |  |  |
|  | 6ga9 | 390fs | 1.8 Å |  |  |
|  | 6gaa | 430fs | 1.8 Å |  |  |
|  | 6gab | 460fs | 1.8 Å |  |  |
|  | 6gac | 490fs | 1.8 Å |  |  |
|  | 6gad | 530fs | 1.8 Å |  |  |
|  | 6gae | 560fs | 1.8 Å |  |  |
|  | 6gaf | 590fs | 1.8 Å |  |  |
|  | 6gag | 630fs | 1.8 Å |  |  |
|  | 6gah | 680fs | 1.8 Å |  |  |
|  | 6gai | 740fs | 1.8 Å |  |  |
|  | 6ga4 | 1ps | 1.8 Å |  |  |
|  | 6ga5 | 3ps | 1.9 Å |  |  |
|  | 6ga6 | 10ps | 1.8 Å |  |  |

**Table S2.** Refinement statistics

| Table S2. Refinement statistics |  |  |  |  |  |
| --- | --- | --- | --- | --- | --- |
| Intermediate | bR | I' | I | J' | J |
| Time period | 0- | < 50 fs | 40-700 fs | 0.5-2 ps | 1-30 ps |
| *C*<sub>10</sub> | 0 | 0 | 3,300 | 0 | -4,200 |
| Coefficient *C*<sub>14</sub> | 0 | 2,000 | 2,700 | 2,700 | 2,000 |
| *C*<sub>17</sub> | 0 | 3,000 | 0 | -1,300 | -300 |
| Starting model | PDB 6g7h |  |  |  |  |
| Resolution range | 50-2.1 Å |  |  |  |  |
| Space group | P1 |  |  |  |  |
| Unit cell | $a = b = 62.32 \text{ Å}; c = 111.10 \text{ Å}; \alpha = \beta = 90^\circ; \text{ and } \gamma = 120^\circ$ | | | | |
| Unique reflections | 80,354 in working set + 4,236 in test set = 84,590 total |  |  |  |  |
| Completeness | 95% in working set + 5% in test set = 100% reconstituted |  |  |  |  |
| *R* (%) | 1.8 | 29.4 | 31.0 | 29.1 | 30.0 |
| *R*<sub>free</sub> (%) | 1.9 | 31.1 | 32.4 | 30.4 | 30.7 |
| Refined content | 230 protein residues + retinal + water molecules |  |  |  |  |
| Number of atoms | 1,798 | 1,795 | 1,798 | 1,796 | 1,795 |
| Water molecules | 8 | 5 | 8 | 6 | 5 |
| RMSD bonds (Å) | 0.005 | 0.009 | 0.009 | 0.009 | 0.009 |
| RMSD angles (°) | 0.793 | 1.206 | 1.105 | 1.085 | 1.068 |
| Rama. favored (%) | 98.7 | 96.5 | 95.6 | 96.1 | 96.5 |
| Rama. outliers (%) | 0.0 | 0.0 | 0.4 | 0.4 | 0.4 |
| Clash score | 4 | 9 | 5 | 4 | 6 |

## Supplementary Figures and Legends

**Figure S1.**
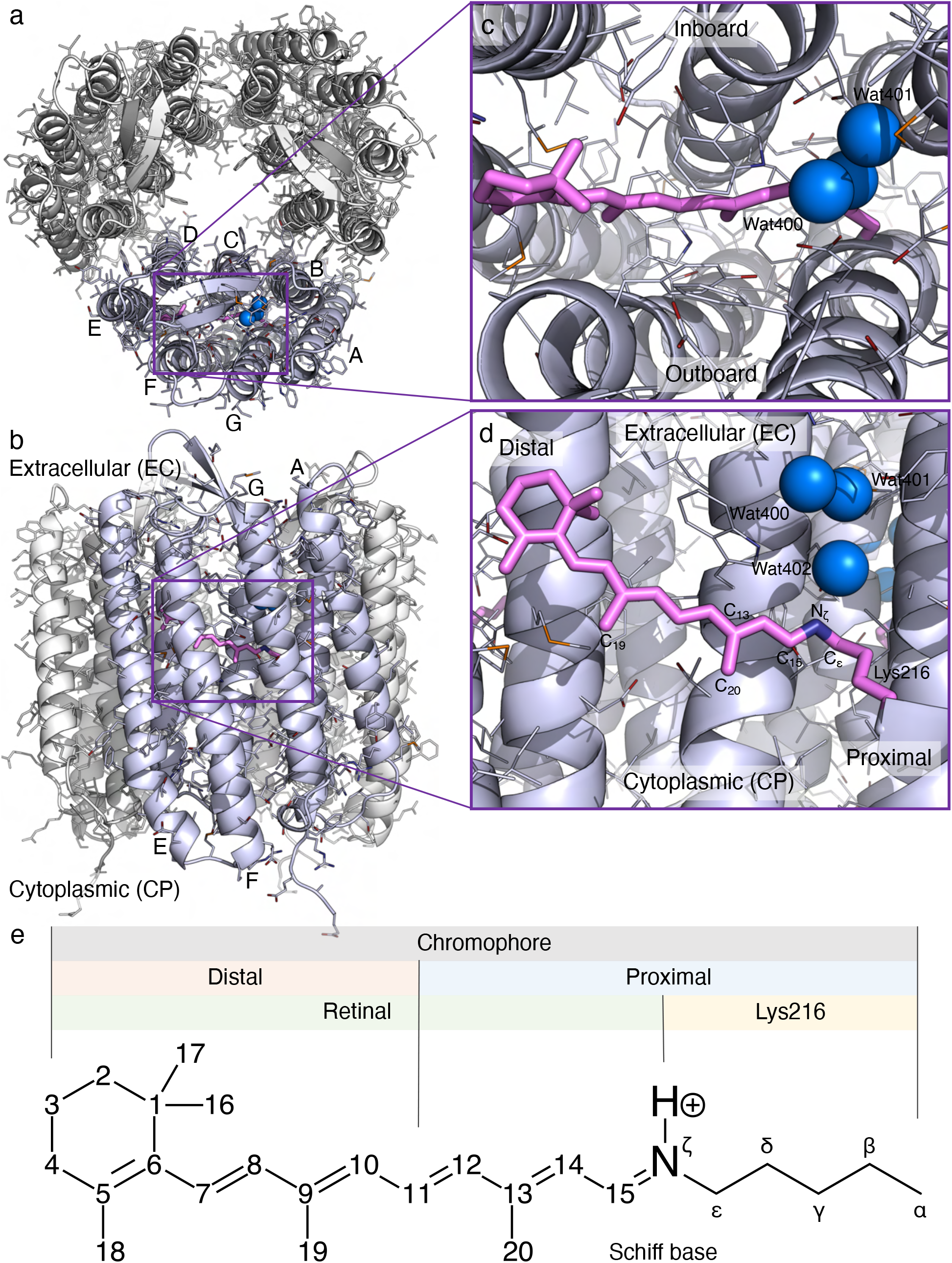
Orientations in bacteriorhodopsin. (a) Bacteriorhodopsin (bR) trimer viewed from the extracellular (EC) side along the three-fold axis. (b) An orthographical view to (a) looking from the outside of the trimer. (c and d) Two orthographical views of the retinal chromophore looking along the three-fold and normal to the three-fold axis. The plane of retinal is largely parallel to the three-fold axis. Therefore, two sides of the plane are called inboard and outboard with respect to the three-fold axis. The direction toward the anchor Lys216 is called proximal. The β-ionone ring direction is therefore distal. (e) Chemical structure of retinal incorporated to its anchor Lys216. The atom numbers and various segment names are marked.

**Figure S2.**
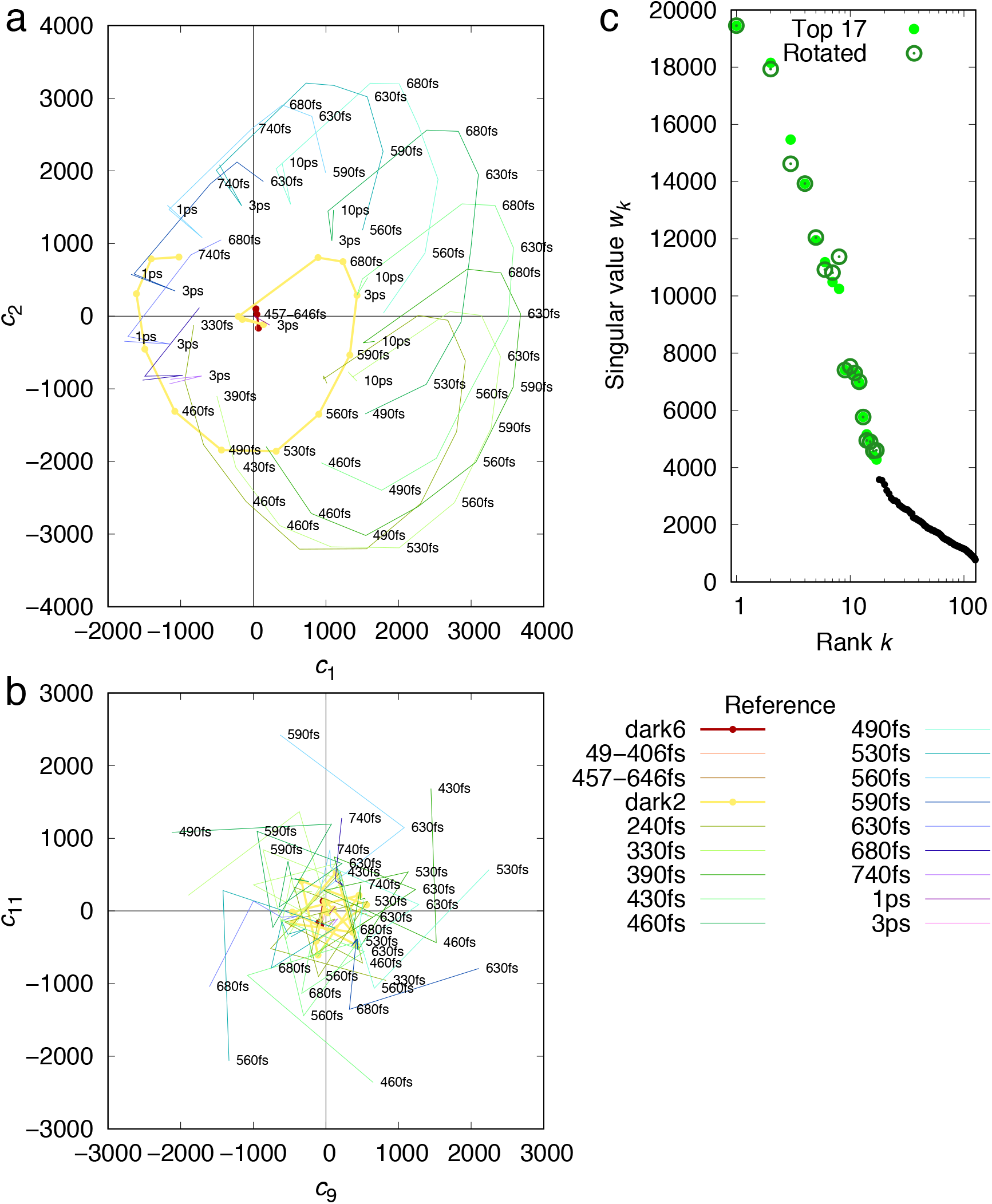
SVD applied to difference Fourier maps. Difference Fourier maps at the short delays *t* ≤ 10 ps are decomposed into component maps. Each difference map at a time delay *t* can be represented by a linear combination of these components, *c*_1_(*t*)***U***_1_ + *c*_2_(*t*)***U***_2_ + …, where ***U***_*k*_ are the time-independent components and *c_k_*(*t*) are their corresponding time-dependent coefficients (Methods). (a and b) Two example plots show circular correlations between *c*_1_ and *c*_2_, *c*_9_ and *c*_11_. These circular correlations indicate two-dimensional oscillations. Each colored trace represents difference maps in a time series calculated with a common reference. Those time series with a dark reference are plotted with thick lines. Other series are in thin lines. (c) Singular values before and after Ren rotation (Ren, 2016, 2019) (Methods). Singular values derived from SVD indicate the significance of the components. 17 of them stand out.

**Figure S3.**
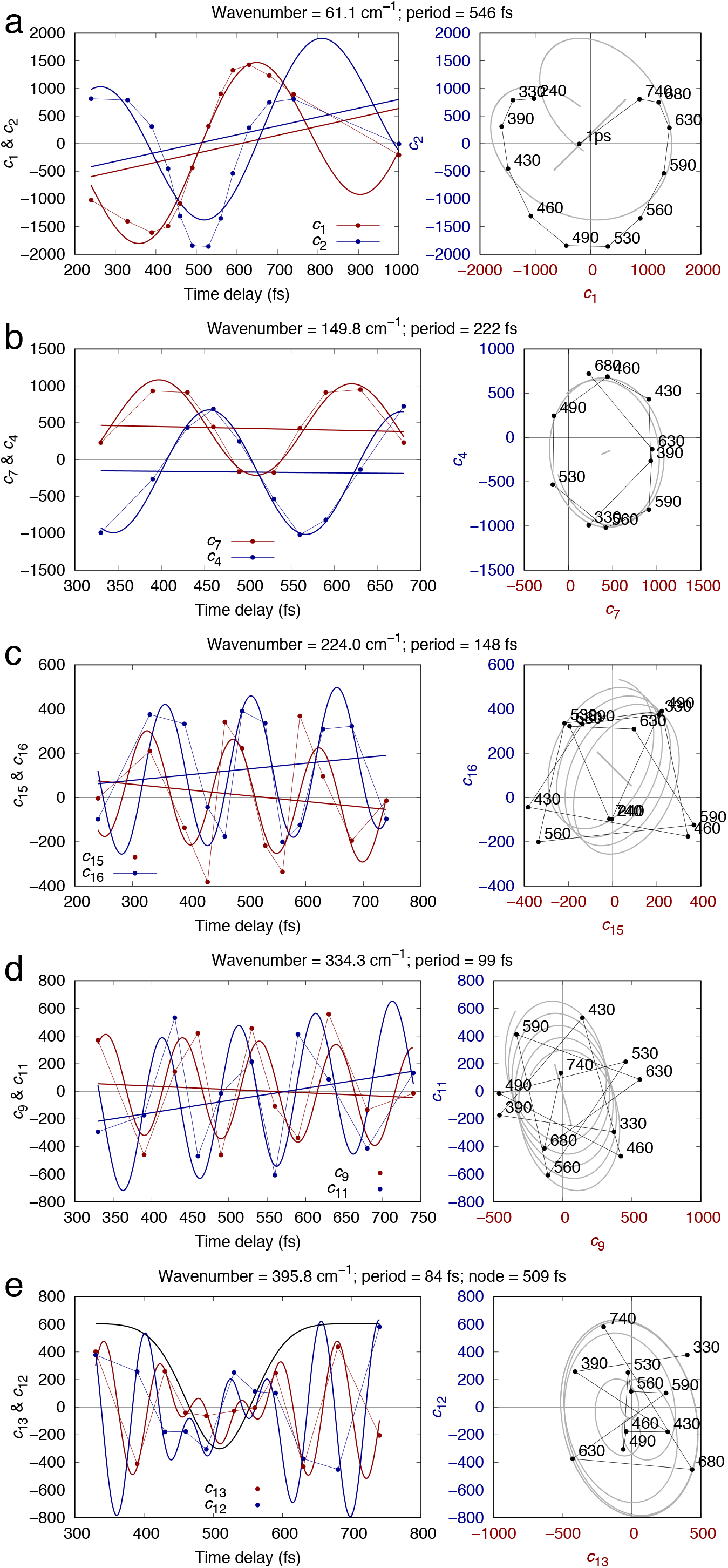
Oscillations of SVD components. The coefficients of ten components *c*_1_, *c*_2_; *c*_4_, *c*_7_; *c*_15_, *c*_16_; *c*_9_, *c*_11_; and *c*_12_, *c*_13_ are found oscillating at frequencies ranging from 60 to 400 cm^-1^. Each pair of the coefficients oscillate at a common frequency. These frequencies are 61±2, 150±3, 224±7, 334±8, and 396±3 cm^-1^, respectively. These coefficients are plotted against the time delay *t* (left) and against each other in a pair (right). Each coefficient is fitted with a sine function around a straight baseline 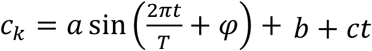. Both the fitted function and the baseline are plotted. The amplitude *a* for the last pair of coefficients *c*_12_ and *c*_13_ are replaced with a Gaussian function 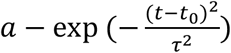 to implement a node at *t*_0_ = 509±5 fs (e).

**Figure S4.**
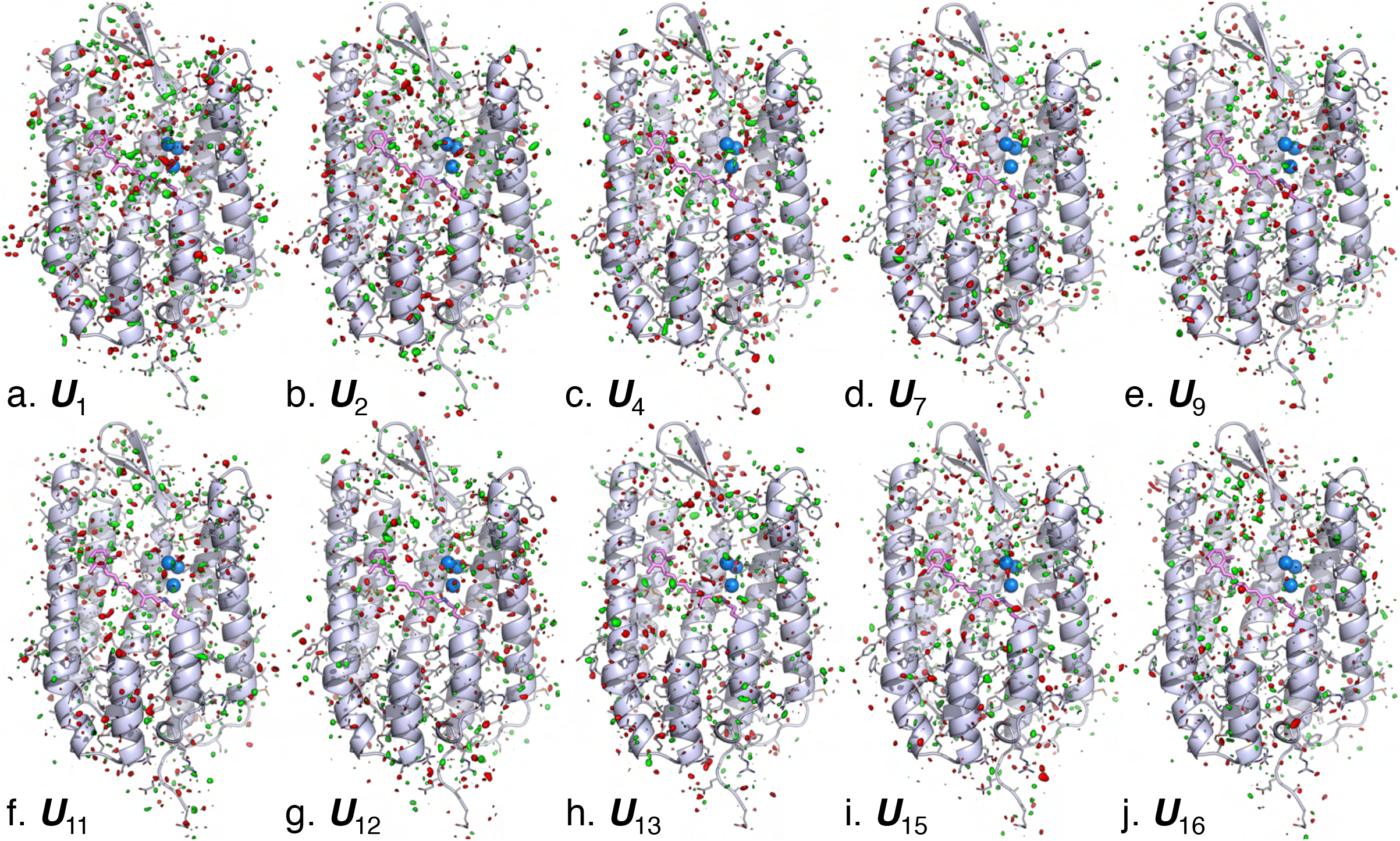
No structural signal in oscillating components. Ten oscillating components are contoured at ±3*σ* in green and red, respectively. The main chain and side chains of the protein are rendered with ribbon and sticks, respectively. The retinal and Lys216 are in purple sticks. Several key waters are in blue spheres. Parts of the structure are omitted to reveal more of the interior. Despite that the time-dependent coefficients to these components contain strong oscillatory signals (Figs. S2 and S3), these components themselves display no obvious association with any structural features such as the retinal or secondary structures. They are in stark contrast to the signal distributions of the non-oscillating components (Figs. 2ab, 3c, and S5).

**Figure S5.**
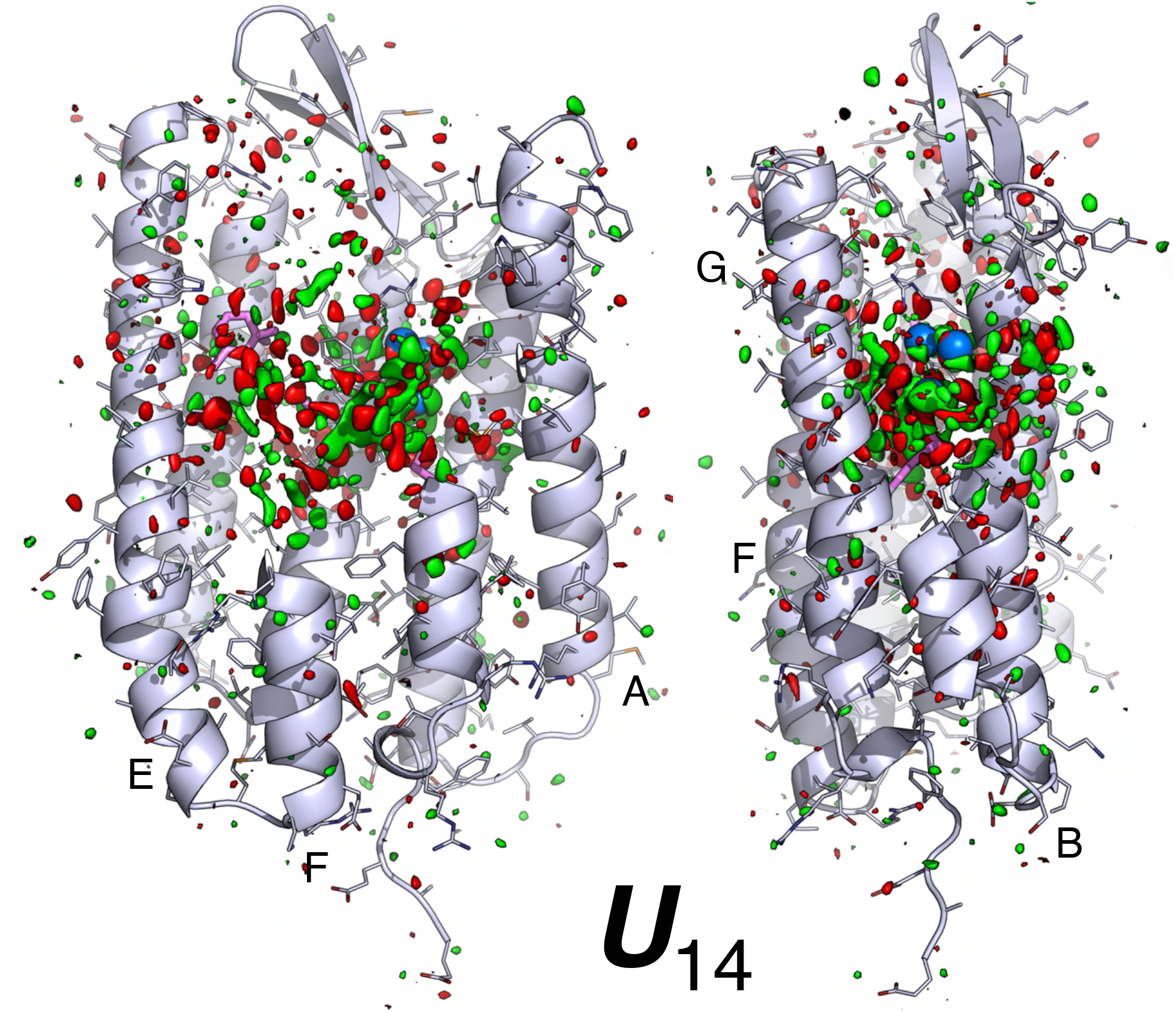
Two orthographical views of component map ***U***_14_. The main chain and side chains of the protein are rendered with ribbons and sticks, respectively. The retinal and Lys216 are in purple sticks. Several key waters are in blue spheres. Parts of the structure are omitted to reveal more of the interior. The map is contoured at ±3*σ* in green and red, respectively. The signals are largely associated with the chromophore and its immediate vicinity.

**Figure S6.**
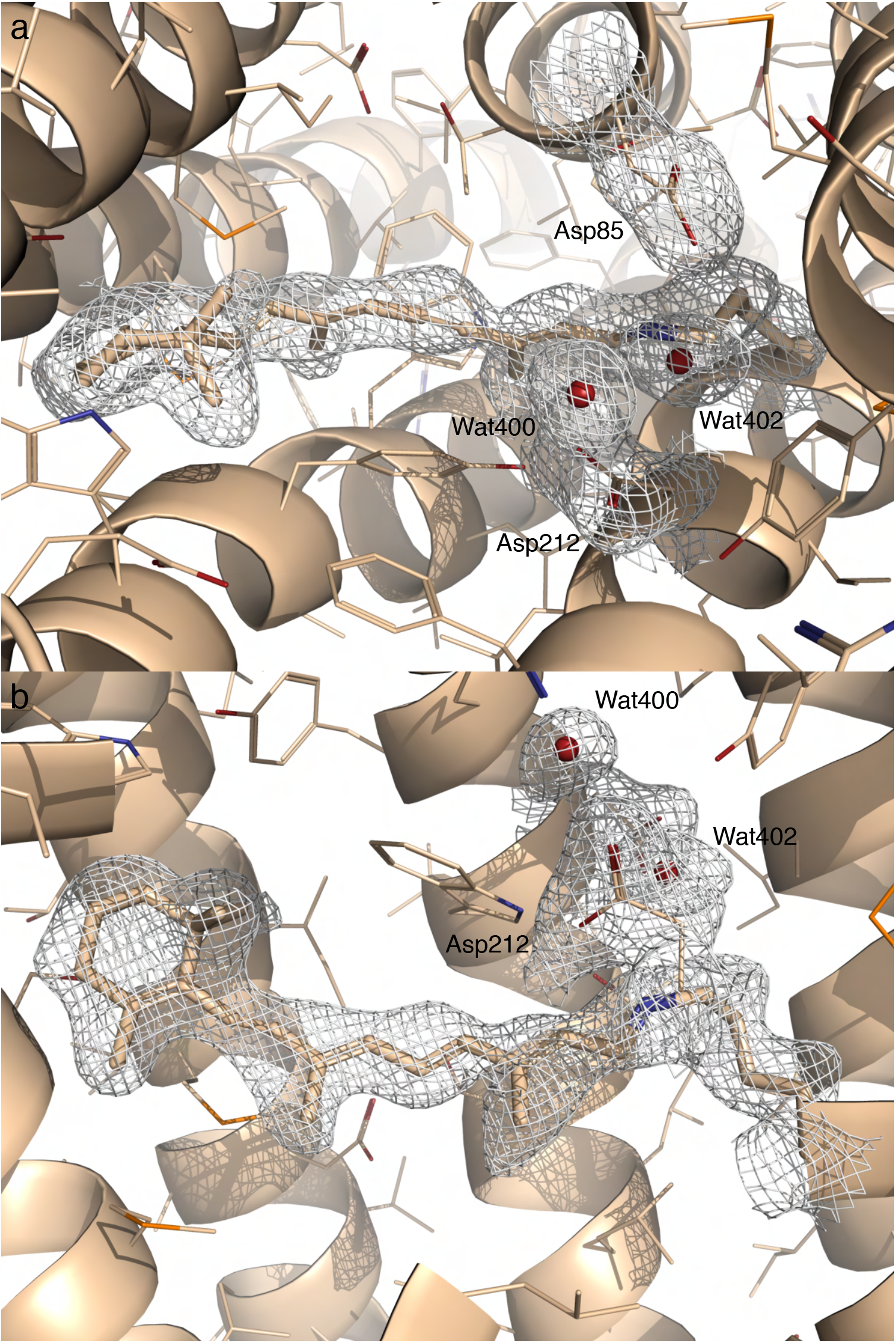
Two orthographical views of the 2Fo-Fc map of I’ contoured at 3.5*σ* Here Fo is the reconstituted structure factor amplitudes rather than observed amplitudes (Table S2). Fc is the structure factor amplitudes calculated from the refine structure (Methods). The same protocol applies to the Fourier synthesis of 2Fo-Fc maps of other intermediates (Figs. S7, S11, and S12).

**Figure S7.**
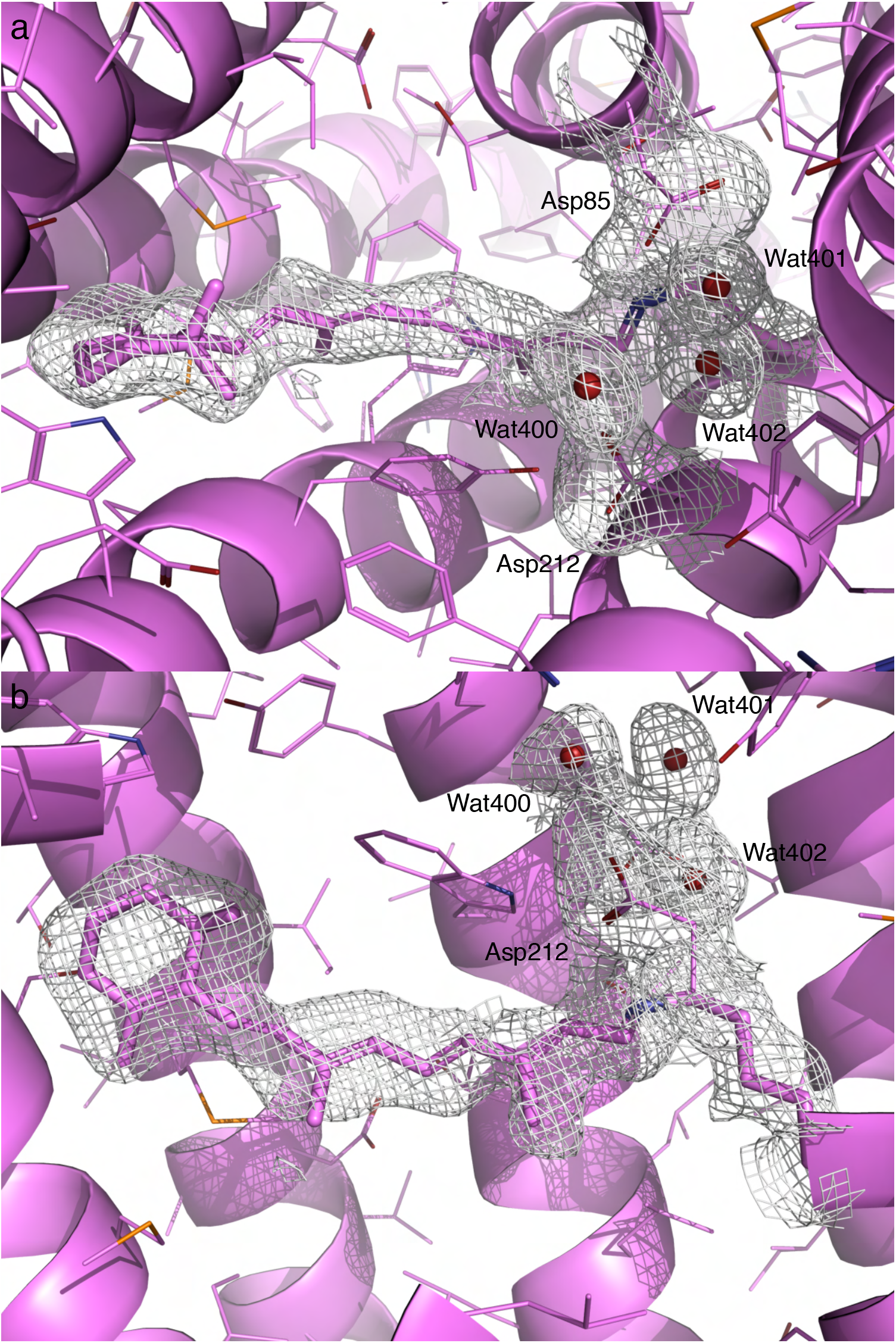
Two orthographical views of the 2Fo-Fc map of I contoured at 3*σ*. Here Fo is the reconstituted structure factor amplitudes rather than observed amplitudes (Table S2). Fc is the structure factor amplitudes calculated from the refine structure (Methods).

**Figure S8.**
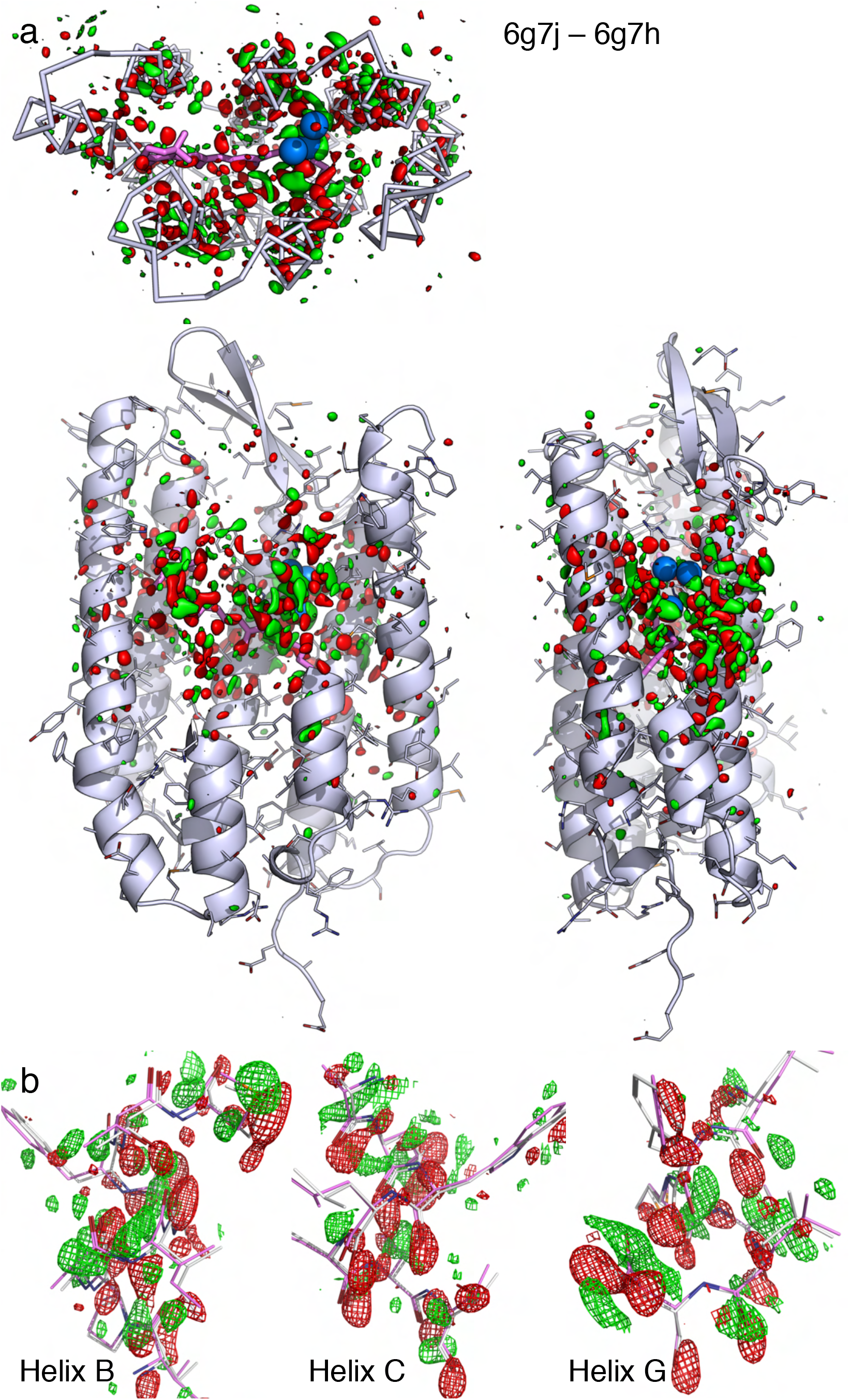
Raw difference Fourier map at 457-646 fs. This difference Fourier map is calculated from the dataset 6g7j at the time point of 457-646 fs by subtracting the dark dataset 6g7h. The map is contoured at ±3*σ* in green and red, respectively. This map is prior to SVD analysis. Compared with ***U***_10_ (Fig. 3c) and the reconstituted map (Fig. 3d), it is clear that this is the original source of the widespread signals except that the *σ* value of this map is higher than those after SVD. (a) The raw difference map contoured in the entire molecule shows the association of the signals with the structural elements at an excellent signal-to-noise ratio. (b) Details of the raw difference map show displacements of helices. The raw difference map is largely the same as the reconstituted map (Fig. 3d).

**Figure S9.**
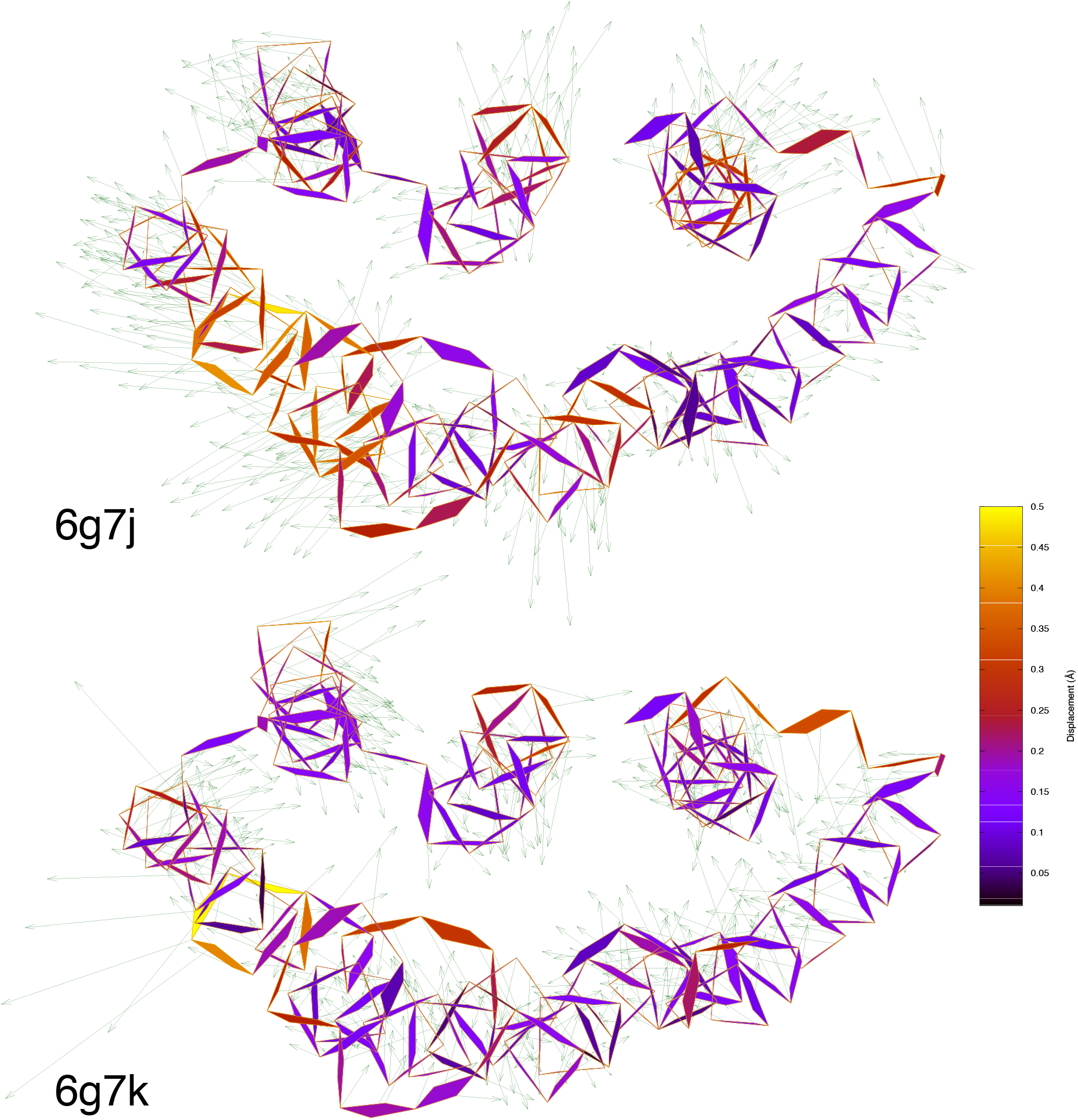
Refined structures at 457-646 fs and 10 ps compared to the ground state. The refined structures of 6g7j and 6g7k and compared with the resting state 6g7h viewed along the trimer three-fold axis from the extracellular side. Atomic displacements in the main chain from bR to 6g7j and 6g7k are color coded and marked by arrows with lengths 20× of the actual displacements. Compared to the structures of I and J species (Fig. 3e), smaller expansion and slight contraction were also captured.

**Figure S10.**
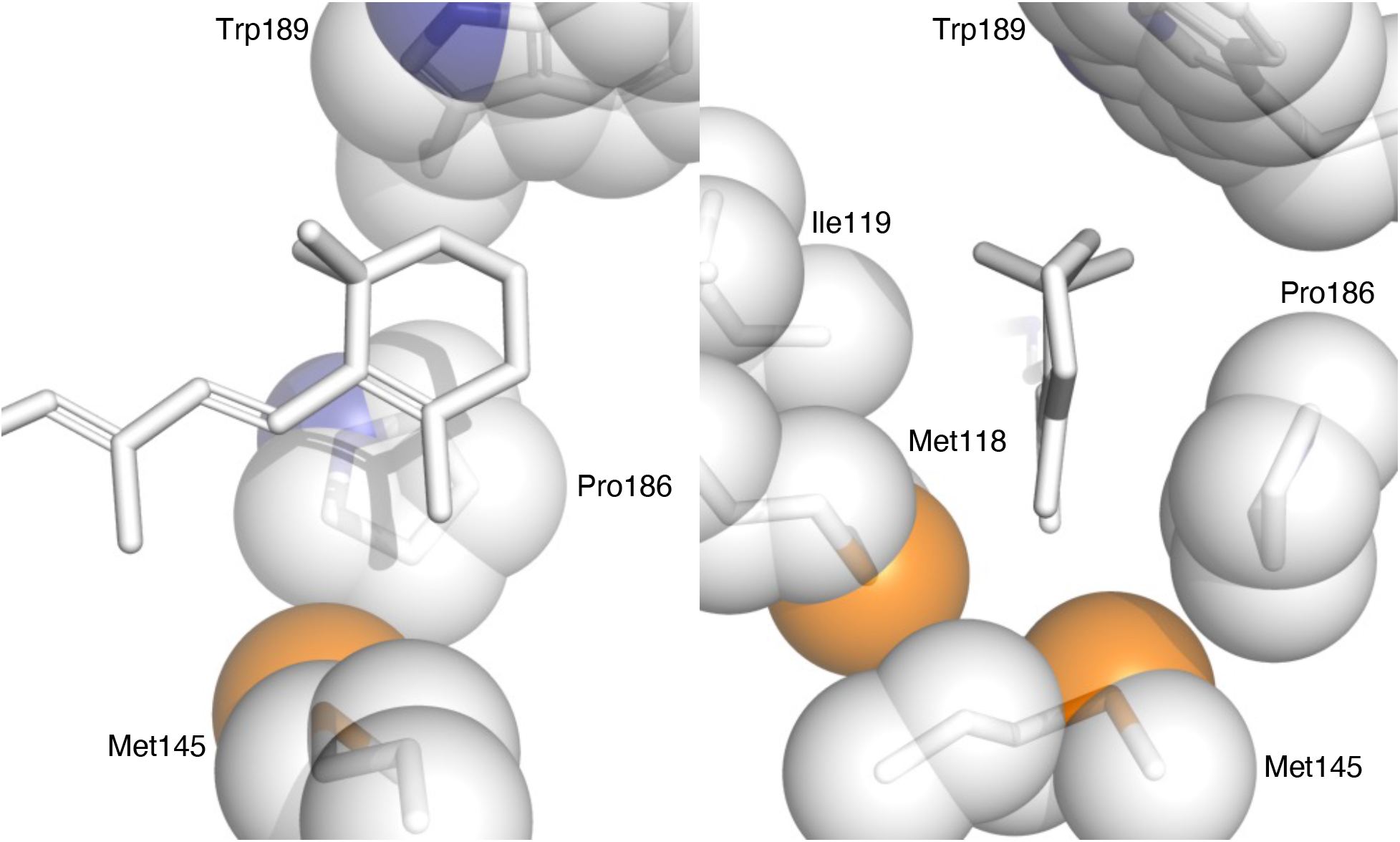
Retinal binding pocket at the distal end. Two orthographical views of the retinal at its distal end. The closest contacts are 3.7 Å to Ile119 and Pro186.

**Figure S11.**
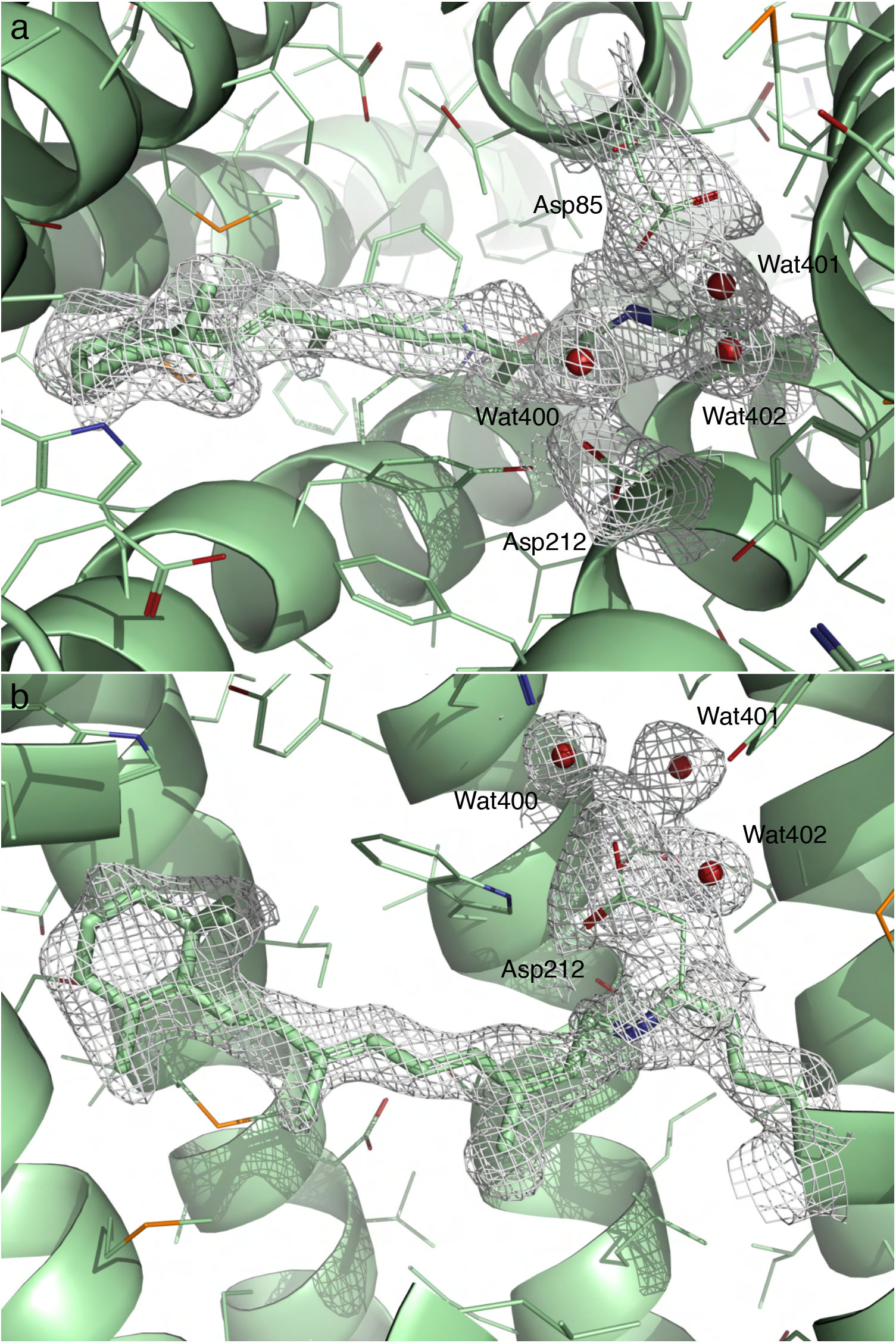
Two orthographical views of the 2Fo-Fc map of J’ contoured at 4*σ*. Here Fo is the reconstituted structure factor amplitudes rather than observed amplitudes (Table S2). Fc is the structure factor amplitudes calculated from the refine structure (Methods).

**Figure S12.**
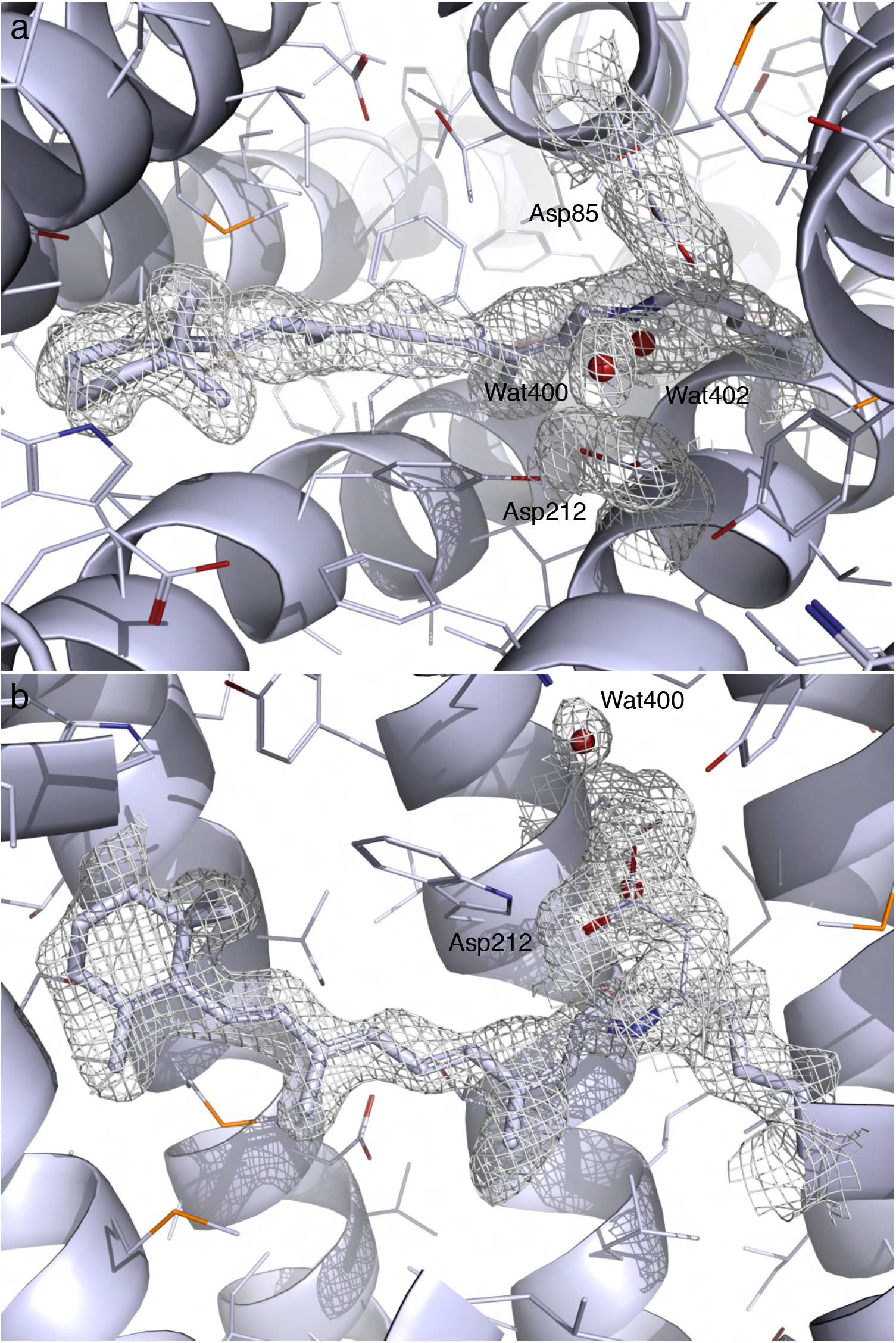
Two orthographical views of the 2Fo-Fc map of J contoured at 5*σ*. Here Fo is the reconstituted structure factor amplitudes rather than observed amplitudes (Table S2). Fc is the structure factor amplitudes calculated from the refine structure (Methods).

## Point-by-point response to peer review

I am grateful to the editor and reviewers for their constructive comments and suggestions. I have clarified a few points and expanded discussion throughout. I have also added several figures for structural comparison as requested. Here I respond to the reviewers’ comments point by point. These comments are copied below in *italics* followed by my response. The topics are arranged by relevance to one another.

*Editor: All three reviewers find the work to be interesting and note that, as written, will require a major revision of our understanding of the biophysics of light induced isomerization of retinal in bacteriorhodopsin.*
*Reviewer 1: The manuscript reports SDV analysis of time-resolved x-ray data directed at the I to J intermediates of the bacteriorhodopsin mechanism. The SDV approach that has been developed over the past decade by the author helps to eliminate noise and systematic error components in the data sets to generate better models of the intermediates. A key new aspect of the study from prior directed at bacteriorhodopsin is the inclusion of structural heterogeneity, providing a truer description of proteins than by modeling based on pure species. The work reveals that excitation-induced charge separation causes retinol to initially form an S shaped conformation with isomerization sampled at many carbons before being fixed at 13C; rapid protein expansion followed by contraction during the retinol motions are uncovered. While bacteriorhodopsin has been extensively studied as a classical model protein, this work more rigorously elucidates structural evolution that provides a more complete explanation for the selectivity of isomerization at the 13C of retinol. The demonstrated functional role of the mechanical flexibility involving such rapid (fs) protein motion is not well appreciated, and I consider this a particularly novel contribution to our general understanding of protein biophysics.*
*Reviewer 3: The paper addresses the mechanism of light induced isomerization of retinal in microbial bacteriorhodopsin in light of recent time-resolved serial femtosecond X-ray crystallography (TR-SFX) experiments. The significance of the work is that photoisomerization of retinal occurs in the case of microbial rhodopsins as well as light reception and signaling in animal rhodopsins. However, at the atomistic level much remains to be learned about the regiospecificity and the stereospecificity and how it is controlled by the protein environment. That knowledge in turn can build a better understanding of light energy conservation, as well as applications to optogenetics as well as understanding visual signaling. The work is potentially appropriate for publication; however, further justification and clarification of the major findings is needed. The paper has some nice insights and commentary which might form the basis for a contribution after it is revised and reviewed again. The research involves a refinement of the previously published time-resolved X-ray structures of bacteriorhodopsin in the range from zero- 10 ps reported by Nogly, Neutze, Scherlter et al. in two earlier papers published in Science. As well there is as a more comprehensive and definitive study by G. Nass Kovacs, I. Schlichting et al. published in Nature Communications (2019). In those papers the results of time-resolved serial crystallography (TR-SFX) were interpreted in terms of photointermediates evolving with time. However, the electron density maps correspond to a mixture of photoexcited species present in the early photocycle. The current paper is a refinement of the previous time-resolved data sets involving four intermediate structures which are refined against the predicted structure factor amplitudes. The method involves singular value decomposition which is clearly and nicely explained in the supplementary material.*

I appreciate the encouraging words of the editor and reviewers as well as their constructive comments.

*Reviewer 3: General Comments: The paper is very well-written with informative and well- prepared figures. The results of the structural refinement involve deconvolution of several distinct early photoproducts which improves the quality of the electron density maps. It is shown that the all-trans retinal undergoes isomerization sampling at the various double bond positions, which are however constrained by the protein binding cavity leading to conformational selection of the* 13-*cis photoproduct. Because of the isomerization sampling, it is proposed that there is a transient expansion of the retinal binding pocket followed by a contraction or a recoil as the 13-cis photoproduct is formed. At this stage, the work is basically a refinement of what is already known.*

The transient, sub-picosecond expansion of the retinal binding pocket and the later recoil that results in a contraction of the binding pocket were not known previously. Such overall response of the protein framework reflects that numerous reaction dead ends fail in simultaneous attempts to isomerize or rotate along the polyene chain of the retinal. The raw difference Fourier map at the key time point of 457-646 fs (Nogly et al., 2018) without the treatment of SVD already shows excellent signal of the pocket expansion (Fig. S8). The refined structures of 6g7j and 6g7k also show the expansion, although to a lesser extent, and a slight contraction (Fig. S9). The previous failure to discern the pocket expansion could be caused by the simplest reason – didn’t look. An alternative explanation could be a suboptimal numerical algorithm or software implementation in calculation or rendering of difference Fourier maps. I have long demonstrated that using difference Fourier maps to reveal weak signal of structural changes is highly nontrivial (Ren et al., 2001). However, these early work of mine received little attention.

*Reviewer 3: The general reader will want to know whether the work is a refinement of the earlier papers, or whether it is transformative with new biophysical insights. My recommendation is that the paper is potential suitable for publication, but that it needs to more fully address all of the points above and below to justify the significance. Are the structures a refinement of the earlier structures or do they lead to transformative insights?*

Whether this work is transformative or merely an incremental improvement from the published literature is up to the readers’ judgement. There is no need to make any claim. However, I have made changes and additions throughout to address every point raised by the reviewers. A major addition is the structural comparison with the previous structures as requested (see below).

*Reviewer 3: However, there is also a proposed mechanism involving a nonspecific Coulombic attraction of the all-trans polyene whose projected length is less in* 13-*cis conformer. The author needs to bring in more persuasively what is known of the excited state photochemistry and molecular orbitals of bacteriorhodopsin, as extensively investigated in earlier work by Birge, El Sayed, Hudson, and others in the earlier literature. The proposal that a greater Coulomb electrostatic force is consistent with the reduced projected length in the* 13-*cis photoproduct as a general mechanism ignores other aspects of the excited state photochemistry. The excited state of a polyene is expected to be more polar consistent with the experimentally measured dipole moment of Mathies et al.*

I appreciate this constructive suggestion, which would no doubt elevate this manuscript to a new level. A brief mention of the barrierless excited state potential energy surface is added (Line 70-73). A number of quantum mechanics and molecular dynamics simulations are discussed and compared to the findings in this study (Altoè et al., 2010; Hayashi et al., 2003). Questions regarding the correspondence between isomerization theory and experimental observation may not have concrete answers yet. At least, I attempt to pose the questions and speculate the correspondence (Line 373-380, 407-420, 433-442, and 476-480).

*Reviewer 3: In addition, one needs to consider the greater anti-bonding character of the excited state molecular orbitals, which is expected to favor an opposite retinal elongation as in the all-trans configuration. There may be other contributions besides the electrostatic and steric contributions described by the author. The resultant energies may not be simply attributable to the light-induced charge separation known from previous experimental work. Probably the stereoselectivity of the protein pocket and conformational selection also come into play, in addition to the Coulombic attractive forces in the isomerization sampling.*
*Reviewer 3: Please clarify whether the comments pertain to the excited singlet state or the ground state potential energy surface. The Coulombic attraction explanation would not work for 11-ciss to all-trans isomerization in visual rhodopsin. Typically, electronic excited states are more polar so one would want to know whether the proposal it unique to rhodopsins or more general. The Coulombic attraction is counteracted by greater anti-bonding character of the excited state molecular orbitals so please explain further. What is the dominant effect, is it the greater dipole moment or the greater anti-bonding character?*

I have been speculating the correspondence between isomerization theories and my structural observation (see above). However, this study does not provide any evidence to support anti-bonding character of the excited state. Quite the contrary, what we can learn here is how a non-specific force causes a highly selected result. Nevertheless, I do not believe that I have intentionally excluded any contribution from other factors. I would be happy to reword if pointed out.

*Reviewer 3: The idea about nonspecific electrostatic forces leading to collapsing of the all-trans retinal by double bond isomerizations along the polyene with conformational selection by the protein is belied by the 11-cis to all-trans retinal isomerization in visual rhodopsin. That would need to be clarified.*
*Reviewer 3: With regard to animal rhodopsins there is ample evidence for expansion of the protein binding cavity upon 11-cis to all-trans isomerization. However, the retinal elongation is opposite to that proposed for bacteriorhodopsin and other microbial rhodopsins.*

It has been reasoned repeatedly that studies on all-trans → 13-*cis* isomerization in bacteriorhodopsin would shed light into the mechanism of 11-cis → all-trans isomerization in visual rhodopsin. The commonality perhaps exists only at a high level in a rather abstract sense, if any. The reactions from all-trans and to all-trans not necessarily share any common trajectory. This is noted in the Concluding remarks (Line 516-523).

*Reviewer 3: Are the refined structures due to multiphoton excitation coming from hot vibrational levels back to the ground state potential surface? Or do they represent alterative non-biological pathways besides the biologically relevant process?*
*Reviewer 3: Very importantly, the role of multiphoton excitation as discussed by Nass Kovacs and also by R.J.D. Miller et al. Nature Communications (2020) needs to be brought more explicitly into the picture, to fully justify the novelty and significance of the present contribution.*
*Reviewer 3: And importantly, whether the intermediate structures correspond to multiphoton excitations, or involve just the first excited singlet state to ground state potentials.*
*Reviewer 3: References – It is essential to mention the paper of R.J.D. Miller et al. Nature Communications 11, 1240 (2020) and comment on the possibilities of multiphoton excitation as an explanation for the coexisting refined structural intermediates (!).*

These comments and suggestions are important and constructive. The refinement of the intermediate structures in this study is largely driven by the captured signals in datasets of Nogly et al. Datasets from Kovacs et al. contribute zero signal. But these datasets contain significant oscillatory signals. I kept the section on low frequency oscillations as short as possible without discussion on multiphoton absorption because it deviates from the main subject of isomerization sampling. The experimental data of Kovacs and Schlichting et al. contain no biologically relevant signal but only oscillations. The reason was pointed out by Miller et al. – multiphoton absorption (Miller et al., 2020). I agree with Reviewer 3 that it would be responsible to discuss in greater detail on the issue of laser peak power and multiphoton absorption rather than leaving it for the readers to conclude. The section on low frequency oscillations has been expanded (Line 157-162 and 176-188).

*Reviewer 1: The main paper also states that the results were based on only the few data sets of Nogly, et al. This point could be clarified, as the data appear to have 17 significant components which would require more than the few data sets. Do the oscillations dominate the sets of Kovacs, et al., such that they contribute little support for U10?*

Yes. The section on low frequency oscillations has been expanded (see above). Discussion on multiphoton excitation is added (Line 117-119 and 176-188). The datasets of Kovacs et al. require a lot of singular values, that is, components. Unfortunately, they do not contribute any useful signal. Their datasets contain near perfect oscillations from 60 to 400 cm^-1^. These oscillations must occur on the potential energy surfaces of multiphoton excitation.

*Reviewer 1: It’s also surprising that a single protein-based component U10 could sufficiently describe both intermediates I and J; does this reflect the crudeness of the current description of the protein response, as one would expect the protein structural changes involved in the expansion around a cis-retinol and contraction for the trans-retinol would differ somewhat in their nature due to the distinct chromophore conformations. Expanded presentation of the specific analysis of bacteriorhopsin in the SI would help to clarify the limitations of the model generated from the study.*

I and J species contain positive and negative ***U***_10_, respectively. But they also require ***U***_14_. Therefore, strictly speaking, a single component is insufficient to describe both intermediates I and J. The coefficients are listed in Table S2. I have also explicitly stated these coefficients in the main text so that some confusion may be avoided (Line 196-197, 208, and 322-325).

*Reviewer 3: To justify the significance of the work, I would expect to see a much stronger comparison to the results of the Nass Kovacs and Nogly papers, with more definitive statements about what is learned of the photointermediate structures versus the simpler intermediate analysis. The paper needs to present a more direct comparison to the earlier structures to more directly substantiate the isomerization sampling and its role in the binding pocket expansion.*

This is a very constructive suggestion. I left these direct comparisons out in the hope that the readers shall conclude by themselves. I added the direct comparison in text and figures and expanded the discussion (Line 282-284, 328-329, 345-371; Figs. 4bc and S9). The key difference is that Nogly et al. and Kovacs et al. presented a gradual rotation of the double bond C_13_=C_14_ instead of discrete states of *trans* and *cis* concluded in this study. The gradual rotation is a misinterpretation of the gradual shift of two or more discrete populations directly caused by the inability to deconvolute mixed signals. This difference is highly consequential to the mechanistic understanding presented in the companion paper (Ren, 2021).

*Reviewer 1: Maybe a matter of taste, but placement of figures could be reconsidered, as in some ways it didn’t follow the sequence of my mind’s digestion of the results. The collection of panels for Figure 2 is primarily the source of this comment. Figure 2a summarizes the structural evolution, so perhaps placing it before all data would be appropriate. I would find it more useful to have Figure 2C focused on the retinol motions placed with information about retinol-based components U14 and U17 in Figure 1. It also seems that Figure 2b which shows the U10 component would be better associated with the panels in Figure 3.*

This is an excellent suggestion. I reorganized the main figures accordingly.

*Reviewer 3: Figures - The figures are high-quality and quite beautiful; however, it is difficult to make cross comparisons due to the small structural differences both in the main text as well as in the supplemental information. It will be more helpful to the reader if an expanded overlay of the refined intermediate structures is provided, together with comparison in particular to the Nass Kovacs structures.*

I added Fig. 4 mainly for structural comparison (Fig. 4bc). Fig. S9 is also added to show the expansion and contraction captured by Nogly et al. but to a lesser extent. These changes in the retinal binding pocket must have been overlooked by the original authors.

*Reviewer 3: Figures - Please provide a figure showing the overlay of the Nogly and Nass Kovacs photointermediate structures so the reader can directly see the extent of the structural differences brought about by the new refinements. This could be combined with overlay of the photointermediate structures, which are lost in Figure (c).*

Overlay of too many structures could be messy. However, the point is well taken. I have managed to present structural comparison without too many figures and extensive overlay of structures.

*Reviewer 3: With regard to figure 2(c) it is necessary to more clearly describe what is actually presented, both in the figure caption and the main text.*
*Reviewer 3: Figure 2 - In part (c) please state more clearly what exactly is plotted in the various panels, both in the caption as well as in the main text. The color coding is not enough.*

An entire paragraph is inserted to describe the conformational parameters (Line 216-238).

*Reviewer 3: Specific Comments: Page 2 bottom - Please clarify the protonation state of the retinylidene Schiff base (PSB or SB) for the general reader, which is important with regards to the proposed non-specific Columbia attractive force.*

Done.

*Reviewer 1: Figure 3(a) caption: “2ab” should be “2b”.*

Corrected.

*Reviewer 1: The one minor recommendation for improvement I have regards the detail of the presentation of the analysis results. The description of the SVD methodology applied is very thorough explained in a general manner in the SI; however, some details of the analysis in this specific study could be more clearly presented. One example, the main paper states that 24 data sets and 18 time points up to 10 ps were analyzed and referred to Table SI1, while Table SI1 lists only 22 data sets under 10 ps.*

The errors in Table S1 are corrected.

## References

Adams, P.D., Afonine, P.V., Bunkóczi, G., Chen, V.B., Davis, I.W., Echols, N., Headd, J.J., Hung, L.-W., Kapral, G.J., Grosse-Kunstleve, R.W., et al. (2010). PHENIX: a comprehensive Python-based system for macromolecular structure solution. Acta Crystallogr. D Biol. Crystallogr. D66, 213–221. https://doi.org/10.1107/S0907444909052925.

Altoè, P., Cembran, A., Olivucci, M., and Garavelli, M. (2010). Aborted double bicycle-pedal isomerization with hydrogen bond breaking is the primary event of bacteriorhodopsin proton pumping. Proc. Natl. Acad. Sci. 107, 20172–20177. https://doi.org/10.1073/pnas.1007000107.

Ansari, A., Berendzen, J., Bowne, S.F., Frauenfelder, H., Iben, I.E., Sauke, T.B., Shyamsunder, E., and Young, R.D. (1985). Protein states and proteinquakes. Proc. Natl. Acad. Sci. 82, 5000–5004. https://doi.org/10.1073/pnas.82.15.5000.

Berman, H.M., Kleywegt, G.J., Nakamura, H., and Markley, J.L. (2012). The Protein Data Bank at 40: Reflecting on the Past to Prepare for the Future. Structure 20, 391–396. https://doi.org/10.1016/j.str.2012.01.010.

Birge, R.R. (1990). Nature of the primary photochemical events in rhodopsin and bacteriorhodopsin. Biochim. Biophys. Acta BBA – Bioenerg. 1016, 293–327. https://doi.org/10.1016/0005-2728(90)90163-X.

Bonvin, A.M.J.J. (2021). 50 years of PDB: a catalyst in structural biology. Nat. Methods 18, 448–449. https://doi.org/10.1038/s41592-021-01138-y.

Brändén, G., and Neutze, R. (2021). Advances and challenges in time-resolved macromolecular crystallography. Science 373, eaba0954. https://doi.org/10.1126/science.aba0954.

Chandonia, J.-M., and Brenner, S.E. (2006). The impact of structural genomics: expectations and outcomes. Science 311, 347–351. https://doi.org/10.1126/science.1121018.

Diller, R., Maiti, S., Walker, G.C., Cowen, B.R., Pippenger, R., Bogomolni, R.A., and Hochstrasser, R.M. (1995). Femtosecond time-resolved infrared laser study of the J-K transition of bacteriorhodopsin. Chem. Phys. Lett. 241, 109–115. https://doi.org/10.1016/0009-2614(95)00598-X.

Ernst, O.P., Lodowski, D.T., Elstner, M., Hegemann, P., Brown, L.S., and Kandori, H. (2014). Microbial and animal rhodopsins: Structures, functions, and molecular mechanisms. Chem. Rev. 114, 126–163. https://doi.org/10.1021/cr4003769.

Freedman, K.A., and Becker, R.S. (1986). Comparative investigation of the photoisomerization of the protonated and unprotonated n-butylamine Schiff bases of 9-cis-, 11-cis-, 13-cis-, and all-trans-retinals. J. Am. Chem. Soc. 108, 1245–1251. https://doi.org/10.1021/ja00266a020.

Glynn, C., and Rodriguez, J.A. (2019). Data-driven challenges and opportunities in crystallography. Emerg. Top. Life Sci. ETLS20180177. https://doi.org/10.1042/ETLS20180177.

Govindjee, R., Balashov, S.P., and Ebrey, T.G. (1990). Quantum efficiency of the photochemical cycle of bacteriorhodopsin. Biophys. J. 58, 597–608. https://doi.org/10.1016/S0006-3495(90)82403-6.

Gozem, S., Luk, H.L., Schapiro, I., and Olivucci, M. (2017). Theory and simulation of the ultrafast double-bond isomerization of biological chromophores. Chem. Rev. 117, 13502–13565. https://doi.org/10.1021/acs.chemrev.7b00177.

Hackett, N.R., Stern, L.J., Chao, B.H., Kronis, K.A., and Khorana, H.G. (1987). Structure-function studies on bacteriorhodopsin. V. Effects of amino acid substitutions in the putative helix F. J. Biol. Chem. 262, 9277–9284. https://doi.org/10.1016/S0021-9258(18)48077-5.

Hamm, P., Zurek, M., Röschinger, T., Patzelt, H., Oesterhelt, D., and Zinth, W. (1996). Femtosecond spectroscopy of the photoisomerisation of the protonated Schiff base of all-trans retinal. Chem. Phys. Lett. 263, 613–621. https://doi.org/10.1016/S0009-2614(96)01269-9.

Hayashi, S., Tajkhorshid, E., and Schulten, K. (2003). Molecular dynamics simulation of bacteriorhodopsin’s photoisomerization using ab initio forces for the excited chromophore. Biophys. J. 85, 1440–1449. https://doi.org/10.1016/S0006-3495(03)74576-7.

Henry, E.R., and Hofrichter, J. (1992). Singular value decomposition: Application to analysis of experimental data. In Numerical Computer Methods, (Academic Press), pp. 129–192.

Herbst, J. (2002). Femtosecond infrared spectroscopy of bacteriorhodopsin chromophore isomerization. Science 297, 822–825. https://doi.org/10.1126/science.1072144.

Joh, N.H., Min, A., Faham, S., Whitelegge, J.P., Yang, D., Woods, V.L., and Bowie, J.U. (2008). Modest stabilization by most hydrogen-bonded side-chain interactions in membrane proteins. Nature 453, 1266–1270. https://doi.org/10.1038/nature06977.

Johnson, P.J.M., Halpin, A., Morizumi, T., S. Brown, L., I. Prokhorenko, V., P. Ernst, O., and Miller, R.J.D. (2014). The photocycle and ultrafast vibrational dynamics of bacteriorhodopsin in lipid nanodiscs. Phys. Chem. Chem. Phys. 16, 21310–21320. https://doi.org/10.1039/C4CP01826E.

Jung, Y.O., Lee, J.H., Kim, J., Schmidt, M., Moffat, K., Šrajer, V., and Ihee, H. (2013). Volume-conserving trans–cis isomerization pathways in photoactive yellow protein visualized by picosecond X-ray crystallography. Nat. Chem. 5, 212–220. https://doi.org/10.1038/nchem.1565.

Kandori, H. (2015). Ion-pumping microbial rhodopsins. Front. Mol. Biosci. 2. https://doi.org/10.3389/fmolb.2015.00052.

Kandori, H., Kinoshita, N., Yamazaki, Y., Maeda, A., Shichida, Y., Needleman, R., Lanyi, J.K., Bizounok, M., Herzfeld, J., Raap, J., et al. (1999). Structural change of threonine 89 upon photoisomerization in bacteriorhodopsin as revealed by polarized FTIR spectroscopy. Biochemistry 38, 9676–9683. https://doi.org/10.1021/bi990713y.

Kiefer, H.V., Gruber, E., Langeland, J., Kusochek, P.A., Bochenkova, A.V., and Andersen, L.H. (2019). Intrinsic photoisomerization dynamics of protonated Schiff-base retinal. Nat. Commun. 10, 1210. https://doi.org/10.1038/s41467-019-09225-7.

Kobayashi, T., Saito, T., and Ohtani, H. (2001). Real-time spectroscopy of transition states in bacteriorhodopsin during retinal isomerization. Nature 414, 531–534. https://doi.org/10.1038/35107042.

Kovacs, G.N., Colletier, J.-P., Grünbein, M.L., Yang, Y., Stensitzki, T., Batyuk, A., Carbajo, S., Doak, R.B., Ehrenberg, D., Foucar, L., et al. (2019). Three-dimensional view of ultrafast dynamics in photoexcited bacteriorhodopsin. Nat. Commun. 10, 3177. https://doi.org/10.1038/s41467-019-10758-0.

Koyama, Y., Kubo, K., Komori, M., Yasuda, H., and Mukai, Y. (1991). Effect of protonation on the isomerization properties of n-butylamine Schiff base of isomeric retinal as revealed by direct HPLC analyses: Selection of isomerization pathways by retinal proteins. Photochem. Photobiol. 54, 433–443. https://doi.org/10.1111/j.1751-1097.1991.tb02038.x.

Lanyi, J.K., and Schobert, B. (2007). Structural changes in the L photointermediate of bacteriorhodopsin. J. Mol. Biol. 365, 1379–1392. https://doi.org/10.1016/j.jmb.2006.11.016.

Lewis, A. (1978). The molecular mechanism of excitation in visual transduction and bacteriorhodopsin. Proc. Natl. Acad. Sci. 75, 549–553. https://doi.org/10.1073/pnas.75.2.549.

Liebel, M., Schnedermann, C., Bassolino, G., Taylor, G., Watts, A., and Kukura, P. (2014). Direct observation of the coherent nuclear response after the absorption of a photon. Phys. Rev. Lett. 112, 238301. https://doi.org/10.1103/PhysRevLett.112.238301.

Liebschner, D., Afonine, P.V., Baker, M.L., Bunkóczi, G., Chen, V.B., Croll, T.I., Hintze, B., Hung, L.-W., Jain, S., McCoy, A.J., et al. (2019). Macromolecular structure determination using X-rays, neutrons and electrons: recent developments in Phenix. Acta Crystallogr. Sect. Struct. Biol. 75, 861–877. https://doi.org/10.1107/S2059798319011471.

Logunov, S.L., Song, L., and El-Sayed, M.A. (1996). Excited-state dynamics of a protonated retinal Schiff base in solution. J. Phys. Chem. 100, 18586–18591. https://doi.org/10.1021/jp962046d.

Marti, T., Otto, H., Mogi, T., Rösselet, S.J., Heyn, M.P., and Khorana, H.G. (1991). Bacteriorhodopsin mutants containing single substitutions of serine or threonine residues are all active in proton translocation. J. Biol. Chem. 266, 6919–6927. https://doi.org/10.1016/S0021-9258(20)89590-8.

Mathies, R., and Stryer, L. (1976). Retinal has a highly dipolar vertically excited singlet state: implications for vision. Proc. Natl. Acad. Sci. 73, 2169–2173. https://doi.org/10.1073/pnas.73.7.2169.

Mathies, R., Brito Cruz, C., Pollard, W., and Shank, C. (1988). Direct observation of the femtosecond excited-state cis-trans isomerization in bacteriorhodopsin. Science 240, 777–779. https://doi.org/10.1126/science.3363359.

McCarty, C.G. (1970). Chapter 9 syn-anti isomerizations and rearrangements. In The Chemistry of the Carbon-Nitrogen Double Bond, (John Wiley & Sons, Ltd), p. 363.

Miller, R.J.D., Paré-Labrosse, O., Sarracini, A., and Besaw, J.E. (2020). Three-dimensional view of ultrafast dynamics in photoexcited bacteriorhodopsin in the multiphoton regime and biological relevance. Nat. Commun. 11, 1240. https://doi.org/10.1038/s41467-020-14971-0.

Mogi, T., Stern, L.J., Hackett, N.R., and Khorana, H.G. (1987). Bacteriorhodopsin mutants containing single tyrosine to phenylalanine substitutions are all active in proton translocation. Proc. Natl. Acad. Sci. 84, 5595–5599. https://doi.org/10.1073/pnas.84.16.5595.

Nogly, P., Weinert, T., James, D., Carbajo, S., Ozerov, D., Furrer, A., Gashi, D., Borin, V., Skopintsev, P., Jaeger, K., et al. (2018). Retinal isomerization in bacteriorhodopsin captured by a femtosecond x-ray laser. Science 361, eaat0094. https://doi.org/10.1126/science.aat0094.

Perálvarez, A., Barnadas, R., Sabés, M., Querol, E., and Padrós, E. (2001). Thr90 is a key residue of the bacteriorhodopsin proton pumping mechanism. FEBS Lett. 508, 399–402. https://doi.org/10.1016/S0014-5793(01)03080-0.

Ren, Z. (2013a). Reaction trajectory revealed by a joint analysis of Protein Data Bank. PLoS ONE 8, e77141. https://doi.org/10.1371/journal.pone.0077141.

Ren, Z. (2013b). Reverse engineering the cooperative machinery of human hemoglobin. PLoS ONE 8, e77363. https://doi.org/10.1371/journal.pone.0077363.

Ren, Z. (2016). Molecular events during translocation and proofreading extracted from 200 static structures of DNA polymerase. Nucleic Acids Res. 6, 1–13. https://doi.org/10.1093/nar/gkw555.

Ren, Z. (2019). Ultrafast structural changes decomposed from serial crystallographic data. J. Phys. Chem. Lett. 10, 7148–7163. https://doi.org/10.1021/acs.jpclett.9b02375.

Ren, Z. (2021). Directional proton conductance in bacteriorhodopsin is driven by concentration gradient, not affinity gradient. BioRxiv doi:10.1101/2021.10.04.463074. https://doi.org/10.1101/2021.10.04.463074.

Ren, Z., Perman, B., Srajer, V., Teng, T.-Y., Pradervand, C., Bourgeois, D., Schotte, F., Ursby, T., Kort, R., Wulff, M., et al. (2001). A molecular movie at 1.8 Å resolution displays the photocycle of photoactive yellow protein, a eubacterial blue-light receptor, from nanoseconds to seconds. Biochemistry 40, 13788–13801. https://doi.org/10.1021/bi0107142.

Ren, Z., Chan, P.W.Y., Moffat, K., Pai, E.F., Royer, W.E., Šrajer, V., and Yang, X. (2013). Resolution of structural heterogeneity in dynamic crystallography. Acta Cryst D69, 946–959. https://doi.org/10.1107/S0907444913003454.

Schaffer, J.E., Kukshal, V., Miller, J.J., Kitainda, V., and Jez, J.M. (2021). Beyond X-rays: an overview of emerging structural biology methods. Emerg. Top. Life Sci. ETLS20200272. https://doi.org/10.1042/ETLS20200272.

Schenkl, S., Mourik, F. van, Zwan, G. van der, Haacke, S., and Chergui, M. (2005). Probing the ultrafast charge translocation of photoexcited retinal in bacteriorhodopsin. Science 309, 917–920. https://doi.org/10.1126/science.1111482.

Schmidt, M., Rajagopal, S., Ren, Z., and Moffat, K. (2003). Application of singular value decomposition to the analysis of time-resolved macromolecular X-ray data. Biophys. J. 84, 2112–2129. https://doi.org/10.1016/S0006-3495(03)75018-8.

Schmidt, M., Graber, T., Henning, R., and Srajer, V. (2010). Five-dimensional crystallography. Acta Crystallogr. A 66, 198–206. https://doi.org/10.1107/S0108767309054166.

Šrajer, V., Ren, Z., Teng, T.-Y., Schmidt, M., Ursby, T., Bourgeois, D., Pradervand, C., Schildkamp, W., Wulff, M., and Moffat, K. (2001). Protein conformational relaxation and ligand migration in myoglobin: A nanosecond to millisecond molecular movie from time-resolved Laue X-ray diffraction. Biochemistry 40, 13802–13815. https://doi.org/10.1021/bi010715u.

Tahara, S., Kuramochi, H., Takeuchi, S., and Tahara, T. (2019). Protein dynamics preceding photoisomerization of the retinal chromophore in bacteriorhodopsin revealed by deep-UV femtosecond stimulated Raman spectroscopy. J. Phys. Chem. Lett. 10, 5422–5427. https://doi.org/10.1021/acs.jpclett.9b02283.

Tittor, J., and Oesterhelt, D. (1990). The quantum yield of bacteriorhodopsin. FEBS Lett. 263, 269–273. https://doi.org/10.1016/0014-5793(90)81390-A.

Ursby, T., and Bourgeois, D. (1997). Improved estimation of structure-factor difference amplitudes from poorly accurate data. Acta Crystallogr. A 53, 564–575. https://doi.org/10.1107/S0108767397004522.

Wang, Q., Schoenlein, R.W., Peteanu, L.A., Mathies, R.A., and Shank, C.V. (1994). Vibrationally coherent photochemistry in the femtosecond primary event of vision. Science 266, 422–424. https://doi.org/10.1126/science.7939680.

Yang, X., Ren, Z., Kuk, J., and Moffat, K. (2011). Temperature-scan cryocrystallography reveals reaction intermediates in bacteriophytochrome. Nature 479, 428–432. https://doi.org/10.1038/nature10506.

Zhong, Q., Ruhman, S., Ottolenghi, M., Sheves, M., Friedman, N., Atkinson, G.H., and Delaney, J.K. (1996). Reexamining the primary light-induced events in bacteriorhodopsin using a synthetic C13=C14-locked chromophore. J Am Chem Soc 118, 12828–12829. https://doi.org/10.1021/ja961058+.

